# A universal workflow for creation, validation and generalization of detailed neuronal models

**DOI:** 10.1101/2022.12.13.520234

**Authors:** Maria Reva, Christian Rössert, Alexis Arnaudon, Tanguy Damart, Darshan Mandge, Anıl Tuncel, Srikanth Ramaswamy, Henry Markram, Werner Van Geit

## Abstract

Detailed single neuron modeling is widely used to study neuronal functions. While cellular and functional diversity across the mammalian cortex is vast, most of the available computational tools are dedicated to the reproduction of a small set of specific features characteristic of a single neuron. Here, we present a generalized automated workflow for the creation of robust electrical models and illustrate its performance by building cell models for the rat somatosensory cortex (SSCx). Each model is based on a 3D morphological reconstruction and a set of ionic mechanisms specific to the cell type. We use an evolutionary algorithm to optimize passive and active ionic parameters to match the electrophysiological features extracted from whole-cell patch-clamp recordings. To shed light on which parameters are constrained by experimental data and which could be degenerate, we perform a parameter sensitivity analysis. We also validate the optimized models against additional experimental stimuli and assess their generalizability on a population of morphologies with the same morphological type. With this workflow, we generate SSCx neuronal models producing the variability of neuronal responses. Due to its versatility, our workflow can be used to build robust biophysical models of any neuronal type.

## 1 Introduction

Biophysically detailed neuronal models enable *in silico* exploration of the underlying complexity of information processing in a single neuron. These models allow systematic and reversible manipulations of the neuronal properties, which are not necessarily feasible in an experimental setup. Detailed neuronal models therefore provide valuable tools for hypothesis testing and to guide further experiments. For example, such models helped to advance our understanding of the importance of morphology on neuronal excitability [Mainen and Sejnowski, 1996, Vetter et al., 2001, van Elburg and van Ooyen, 2010] and the contribution of specific currents to cell function [Hay et al., 2011, Traub et al., 2003, Poirazi et al., 2003, Segev and London, 2000]. In addition they served as a basis to build neuronal circuits to simulate and study brain activity [Billeh et al., 2020, Markram et al., 2015].

In general, electrical models (e-models) are expected to reproduce experimentally observed electrophysiological behaviors. This can be quantified via a similarity score that can be computed either directly as the difference between experimental and numerical traces [Brookings et al., 2014] or between features extracted from these traces. Since parameters such as ion channel conductances and passive membrane properties are currently not always experimentally measurable, obtaining a model with a good score requires either manual or automatic exploration of the parameter space [Prinz et al., 2003, Günay et al., 2008, Sekulić et al., 2014]. The latter can be achieved through stochastic global parameter optimization using evolutionary algorithms (EA). Simple to parallelize and effective in high dimensions, EAs have gained popularity in the field [Van Geit et al., 2008]. Here we use an indicator-based evolutionary algorithm (IBEA) [Zitzler and Künzli, 2004, Van Geit et al., 2016] with good performance according to benchmarks [Mohácsi et al., 2020]. Since cell models are usually customized through feature extraction and parameter fitting for a specific study or released independently, to our knowledge only a few completely open-sourced and reproducible workflows of model optimization exist [Gouwens et al., 2018]. Here we present a fully integrated, single cell model-building routine based on open-source tools.

Model building can serve to construct a cell model that would represent either a single biological cell or a predefined type of cells [Ascoli et al., 2008, Markram et al., 2004, Tasic et al., 2018]. Although the former approach is common [Gouwens et al., 2018], it has a few limitations. First, while building a neuronal model it could be preferable to constrain its parameters by a rich repertoire of measured neuronal behaviors. For example, for a model that includes a full morphological tree, it would be beneficial to tune this model not only to somatic responses but also take into account dendritic recordings. However, usually it is challenging to collect a battery of voltage recordings from several cellular compartments in the same neuron. This can be overcome by combining dendritic and somatic recordings which were acquired in different cells of the electrical type (e-type). Moreover, the task of building neuronal circuits requires constructing hundreds of thousands of neuronal models that represent the same neuronal type. Optimizing the parameters of such a large number of models would be computationally prohibitive and time-consuming. On the other hand, a canonical model allows the study of properties of a neuronal type and can be used in large circuit building by applying the model to a set of morphologies that constitute the same morphological type (Markram et al. [2015]).

In this work, we developed a workflow for single model creation, which allows us to build canonical neuronal models. With this approach we created 40 models representing 11 e-types in the juvenile rat somatosensory cortex (SSCx). For each cell type, we extracted a set of electrophysiological features which were used to optimize model parameters. Then, each canonical model was applied to a number of morphologies to assess its generalization. In addition, using a layer 5 pyramidal cell (L5PC) model as a use case we performed a dendritic model validation and an analysis of the optimized parameter space. In addition, we present a set of notebooks that allows the reader to follow all the steps of the workflow on the example of the L5PC.

## 2 Results

### 2.1 Single cell model building workflow

Our goal was to develop an open-source workflow for generating robust e-models of neuronal cell types (Fig.1). Our workflow capitalizes on other open-source Python packages, such as BluePyEfe [Blue Brain Project, 2020a] for the extraction of electrophysiological features, BluePyOpt [Van Geit et al., 2016] to perform data-driven model optimization and BluePyMM [Blue Brain Project, 2020b] to generalize the electrical models to large sets of neuronal morphologies. The modular structure of our workflow with configuration files containing information about the model’s parameters, features, and cellular morphology allows for a flexible and multifaceted range of applications.

**Figure 1:**
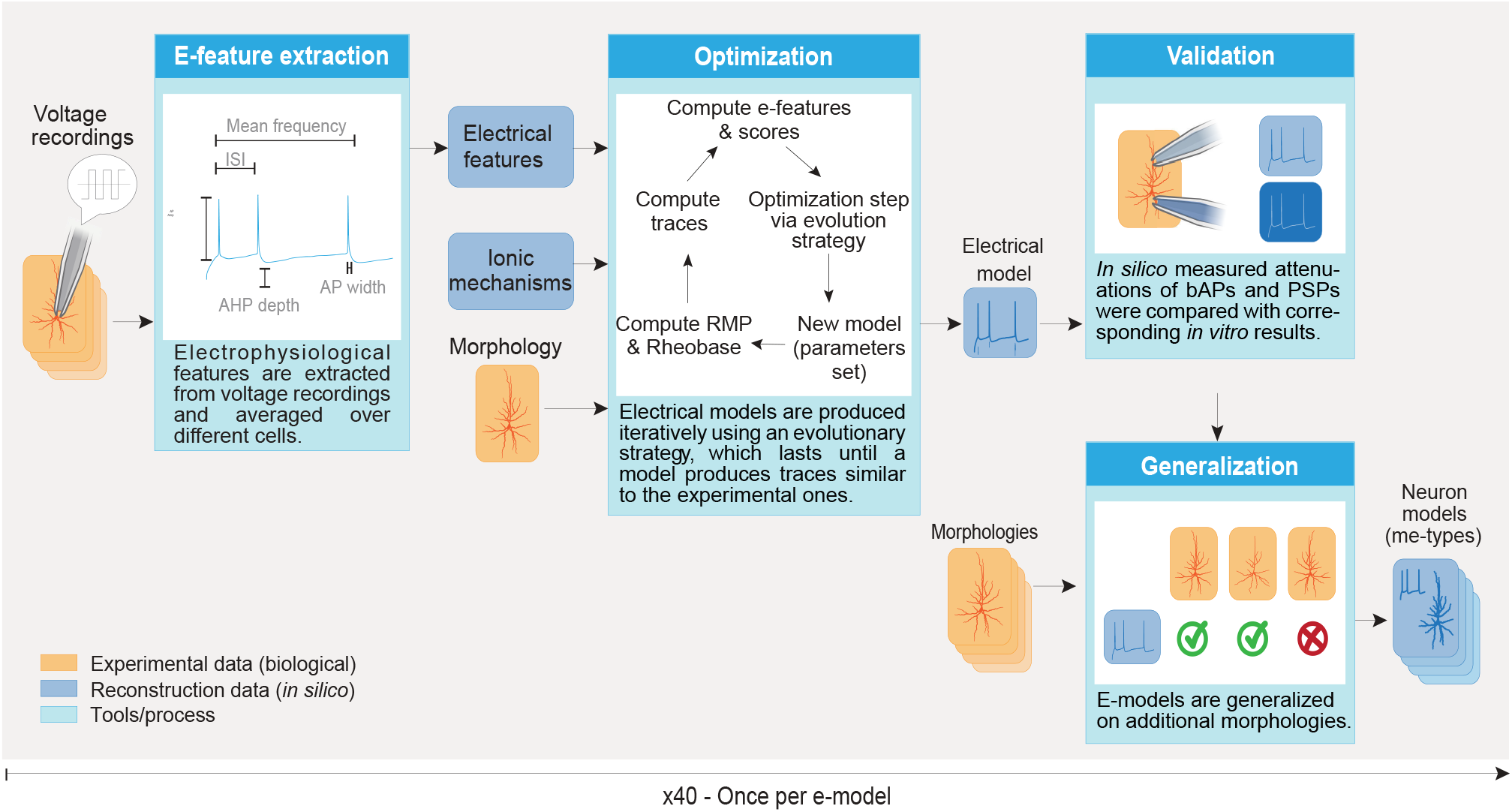
E-model building workflow. Schematic illustration of the steps involved in creating neuronal models. See text of Sect. 2.1 for a description of the pipeline.

During the first step of the workflow we extracted a set of electrophysiological features (efeatures) from the voltage recordings of each cell that belong to the same e-type using BluePyEfe (see Sec.2.2 and Methods). The average value of the extracted features of all the cells of the same e-type were used as constraints during the optimization process (Fig.1, E-feature extraction). The standard deviations were also computed and were used as a way to normalize the e-feature scores during the optimization step.

The second step (Fig.1, Optimization) aims at building canonical e-models that reproduce the previously obtained e-feature values when the neuron model is stimulated *in silico* with the same current stimulus. An e-model is composed of: an detailed morphological reconstruction (exemplar morphology), mechanisms describing the dynamics of the passive and active electric properties of the membrane, and mechanisms describing the ionic dynamics inside the membrane of the cell (see Table 5). The free parameters of these mechanisms (e.g. maximum channel conductance, calcium decay) were optimized using an evolutionary algorithm [Zitzler and Künzli, 2004, Van Geit et al., 2016] with BluePyOpt. The cost function for the optimization is the sum of the errors between e-features produced by the e-model and the experimental e-features (Eq. 1). The evolutionary algorithm then searches for a set of parameters that minimize this cost function.

Once e-models were built, their quality was assessed by inspecting behaviors that were not specifically constrained with experimental data during the optimization (Fig.1, Validation). This step checks for the reproduction of the attenuation of dendritic and synaptic potentials as well as somatic responses to the stimuli not used during optimizations (see Sect.2.4 for details).

Finally, we tested morphological generalizability of the e-models using the BluePyMM software [Blue Brain Project, 2020b]). Since our models were optimized for a single morphology, we examined their performance on a broad collection of morphologies to select working pairs of morphologies and e-models, leading to what is referred to as morpho-electric models (me-models) (Fig.1). This collection of in *silico* cells can for example be used to build a microcircuit Markram et al. [2015] (See Sect.2.5 and Methods sections).

With our workflow, we generated e-models for the various firing and morphological neuronal types present in the juvenile rat SSCx. Based on the whole-cell patch-clamp single-cell recordings each cell was manually assigned to one of 11 e-types [Ascoli et al., 2008]: continuous adapting pyramidal cells (cADpyr) and 10 interneuron firing e-types: continuous accommodating (cAC), burst accommodating (bAC), continuous non-accommodating (cNAC), burst nonaccommodating (bNAC), delayed non-accommodating (dNAC), delayed stuttering (dSTUT), burst irregular firing (bIR), continuous irregular firing (cIR), burst stuttering bSTUT) and continuous stuttering (cSTUT). One e-model was generated per interneuron firing type. For the pyramidal cells one e-model per layer (2/3, 4, 5, 6) was generated. For e-models that did not generalize well to all the matching morphological types (m-types), additional optimizations were run to generate e-models for specific m-types. This resulted in a final set of 40 e-models.

In the next chapters, we present in more details each step of our workflow.

### 2.2 Electrophysiological features of the SSCx neuronal e-types

Using the first step of the workflow, we extracted a number of e-features from voltage traces for each neuron that belongs to one of the 11 e-types identified in our SSCx recordings. To demonstrate variability between different e-types, we considered somatic single-cell recordings in response to a depolarizing step current injection (2 s) with different intensities. The features that were extracted from these recordings represent firing properties of the cell, such as the mean firing frequency, interspike intervals and burst number. Then, a hyperpolarizing current (3 s) was applied to characterize passive cell properties and voltage sag. Finally, a short (50 ms) depolarizing step with a high sampling rate was applied to reliably record a single action potential (AP) waveform (Fig.2A). Consequently, features that were extracted from these recordings target single AP properties, such as afterhyperpolarization potential (AHP), action potential (AP) width, AP fall and rise time.

**Figure 2:**
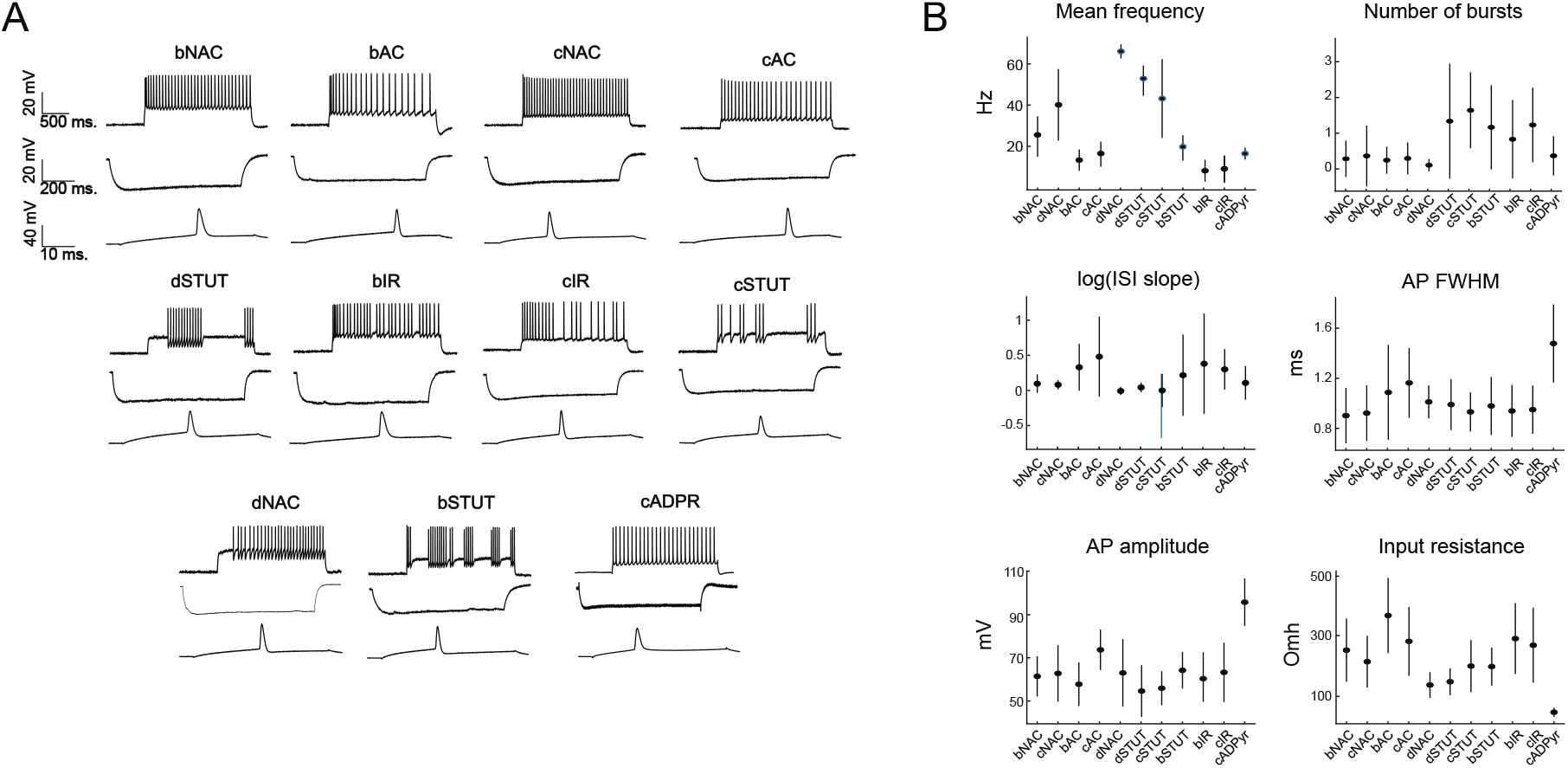
E-features of the SSCx neuronal e-types. A. Exemplar patch-clamp voltage recordings of 11 e-types. Each subplot consists of exemplar traces for three stimuli: long depolarizing current (2 s), hyperpolarizing current (1 s) and a short depolarizing current (50 ms). B. Exemplar subset of extracted features from experimental recordings for each e-type. The efeatures presented here are: firing frequency (for injected current corresponding to 300% from the rheobase), number of bursts (current of 150% from the rheobase), logarithm of the slope of interspike intervals (‘log(ISI slope)’, current of 150% from the rheobase), AP full width at half maximum (‘AP FWHM’, current of 150% from the rheobase), AP amplitude (current of 150% from the rheobase) and input resistance (current of −40 pA). All features are plotted as mean value ± the standard deviation.

To systematically combine e-features from different cells within the same e-type, we deployed the following normalization strategy. First, from the available recordings we computed a threshold current (rheobase) for each cell. This value was used to re-scale all the protocols (see Methods) such that each protocol corresponds to a percentage of the rheobase. Then, for each chosen protocol intensity (e.g. 150% of the rheobase) we selected the corresponding voltage traces and extracted a set of features from each recording. For each e-type, we then computed the mean and standard deviation of the feature values (Fig.2B) which were used in the next step of the workflow.

We illustrate the diversity of neuronal firing patterns considered in the current study for several extracted e-features in Fig.2B (and SI Table 3). As expected, pyramidal cells (i.e. cADpyr e-type) exhibit lower input resistance than inhibitory cells. On average, all inhibitory cells have shorter AP duration and lower AP amplitude when compared to cADpyr cells. Among the inhibitory cells, accommodating and irregularly firing cells have lower firing rates than nonaccommodating and stuttering ones, while stuttering and irregularly firing types have a higher number of bursts than other firing types. Moreover, typically irregularly firing, bursting and accommodating e-types show more diversity in their interspike intervals (ISI), reflected in the logarithmic slope of ISIs, than the rest of the firing e-types. This observation is coherent with the firing pattern of the stuttering and irregularly bursting cells, which can display a wide variability in the number of bursts and interspike intervals.

### 2.3 Model construction and optimization

In order to create an e-model of a neuron, we needed to define ionic mechanisms (see the ion channels and calcium dynamics in Fig.3A), along with previously mentioned experimental constraints (e.g. e-features and reconstructed neuronal morphologies).

**Figure 3:**
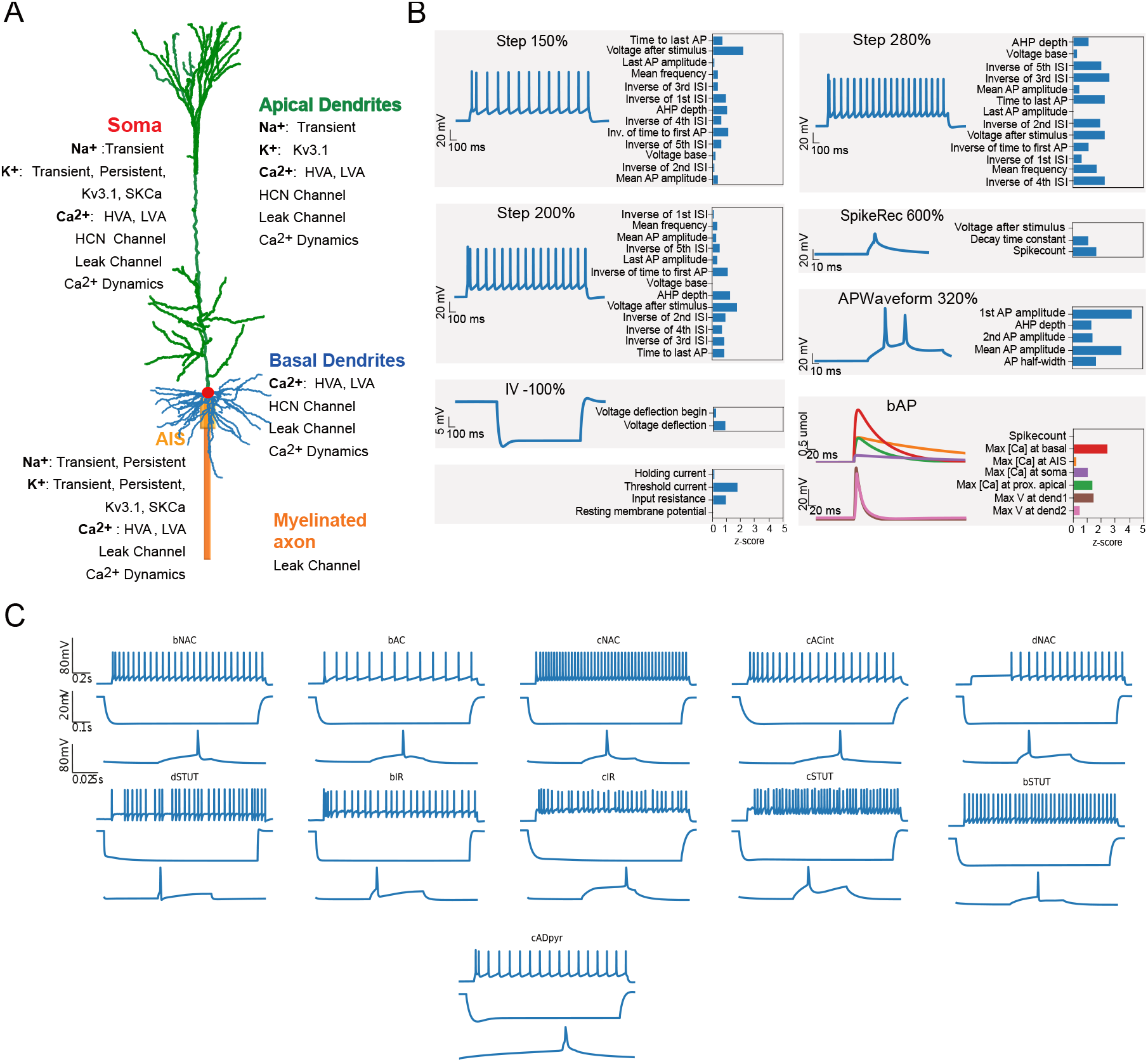
Model construction and optimization results. A. Layer 5 thick-tufted pyramidal cell (L5PC) morphology showing the mechanisms inserted in different morphological sections: the apical dendrites (green), soma (red), basal dendrites (blue), axon initial segment, AIS (thin yellow cylinder) and myelinated axon (thick orange cylinder). B. Scores and exemplar traces of the optimized L5PC model for e-features of single action potentials (APWaveform), firing properties (Step/IDRest), input resistance and hyperpolarization features (IV), back-propagating action potential and peak intracellular calcium concentration recorded in the apical dendrites, soma and AIS, (bAP, for pyramidal neurons only) and spike recovery (SpikeRec). The resting membrane potential (RMP), holding current, threshold current are also optimized as e-features. C. *In silico* voltage recordings obtained from e-models (one for each e-type) for three protocols: IDRest (150/140 % from the rheobase), IV (−100 % from the rheobase) and APWaveform (320/350 % from the rheobase). These e-models are in close agreement with various experimental e-types as shown in Fig.2A

The models of ion channels were constructed based on the Hodgkin-Huxley formalism (see Methods and Fig.7). Each e-model has a number of ionic mechanisms (e.g. persistent and transient sodium, persistent and transient potassium, intracellular calcium dynamics, see Table 5). Several interneuron e-types (bIR, cIR, cSTUT, bSTUT, dSTUT) additionally have a stochastic potassium channel [Diba et al., 2006, Mendonça et al., 2016] for irregular spiking. All aforementioned mechanisms were implemented in the NEURON simulator Carnevale and Hines [2006]. A full list of the ionic mechanisms used in this work can be found in Table 7. While models of these ionic mechanisms were defined with respect to their experimental characteristics, some of their parameters, such as the exact maximal ionic conductances or intracellular calcium decay, remain unknown. The aim of the optimization is to find a set of parameters that will allow the model to accurately reproduce cellular features extracted from the experiments.

To solve this task, we used the indicator-based evolutionary algorithm (IBEA) algorithm [Zitzler and Künzli, 2004] available in the Python package BluePyOpt [Van Geit et al., 2016] (see Methods). At each generation of the evolutionary process, a wide range of models are produced through random mutations and mating of the members of the previous generations. The fitness of the offspring is then evaluated by computing the difference between the e-features produced by the e-model and the experimental ones. The parents of the next generation are selected based on their fitness. Due to the use of random numbers in the algorithm, this optimization process can be repeated several times with different random seeds, until satisfactory models are obtained.

The e-models are evaluated according to a two-step process. First, the resting membrane potential (RMP), input resistance (Rin), holding current, and rheobase of the models are computed using respectively: no stimulus, a small negative stimulus, constant stimuli of increasing amplitude, and step stimuli of increasing amplitude. Second, the e-models are evaluated for a number of step protocols that reveal the firing properties of the neuron. The current stimuli that are used were individually rescaled based on the holding current and rheobase.

The results of the optimizations are illustrated for a thick-tufted pyramidal neuron e-model (cADpyr L5TPC) in Fig.3B. The majority of e-feature scores (z-scores) for this e-model are below two standard deviations of the experimental mean value – suggesting the model closely replicates the experimental data. The e-model was optimized for multiple starting seed values and the best seed was chosen for each e-model.

When optimizing interneuron models that contain stochastic potassium channels we used a two-step optimization. This is due to the fact that the optimization did not converge with all the parameters free at the same time. During the first stage, the stochasticity of the potassium channels was disabled and we optimized for all the e-features except burst number. In the second optimization stage, stochasticity was re-enabled and all the parameter values obtained from the first stage were used, except for the maximum conductance of stochastic potassium channels. This part of the optimization was constrained by burst number and interspike intervals. Such an approach also reduces the optimization time since the non-deterministic mode of stochastic voltage-gated potassium (Kv) channels increases the run time of simulations.

Using this procedure, we optimized e-models for all aforementioned SSCx e-types. Exemplar responses from different e-models representing all the e-types are depicted in Fig.3C. Note the similarity in the firing patterns between the e-models and the experimentally recorded data (Fig. 2A). Once the best e-model is found for an e-type, it undergoes validation.

### 2.4 Validation and analysis of the detailed neuronal model

It was unknown how the optimized e-models would respond to the stimuli not used in the process of optimization. To test this we applied two validation routines to the study case of L5PC: dendro-synaptic (Markram et al. [2015]) and somatic. Since dendritic attenuation data was available only for the L5PC, the validation and e-model analyses were performed only for this e-model.

First, we tested whether our model can reproduce the attenuation of dendritic back-propagating action potentials (bAPs) and excitatory postsynaptic potentials (EPSPs) observed in the literature. For bAP attenuation, we injected a current step (5 ms, 2 nA) in the soma and recorded the voltage from apical and basal dendrites at various distances from the soma (Fig.4A). We compared the attenuation length constant of our *in silico* results to the experimentally reported ones [Berger et al., 2001, Nevian et al., 2007] (Fig.4B). For validation of EPSP attenuation we simulated a transient change in the synaptic conductance in distant apical (1.5 nS) and basal (0.2 nS) dendrites (Fig.4C). Then, we calculated the ratio of dendritic to somatic attenuation and compared it to the experimental results reported previously [Berger et al., 2001, Nevian et al., 2007]. Both bAP and EPSP attenuation in the basal dendrites in the e-model (bAP: 120.6 ± 1.1 *μm*, EPSP: 39.6 ± 0.2 *μm*) were consistent with the experimentally reported values (bAP: 145.8 ± 8.7 *μm*, EPSP: 39.9 ± 0.9 *μm*). In the case of the apical dendrite, agreement between in *silico* (bAP: 651.3± 1.8, EPSP: 263.6± 1.8 *μm*) and experimental (bAP: 675.5.3±27.4 *μm*, EPSP: 273.7 ± 8.2 *μm*) results were achieved for the in *silico* morphologies with apical dendrites with a diameter of 2 — 6 *μm* (Fig. 4B-C). These diameter values are larger than those found in the juvenile rat [Zhu, 2000], which is consistent with the fact that the aforementioned studies were performed in the adult rat.

**Figure 4:**
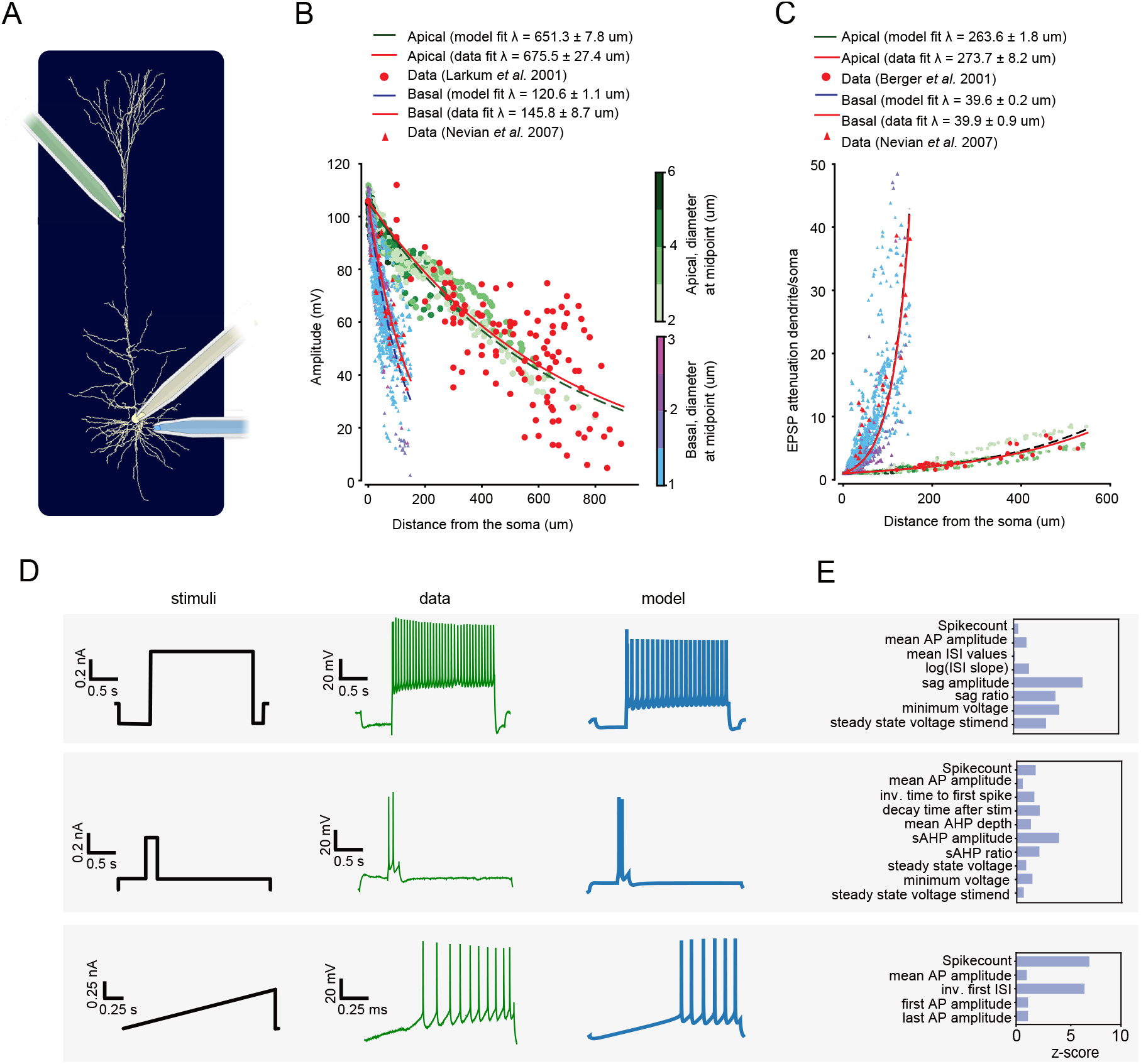
Synaptic and somatic validations of L5TPC electrical model. A. Schematic representation of the experimental setup. B. Predicted bAP amplitude measured at different locations on apical (green dots) or basal dendrites (blue dots) and experimental results from literature (red dots). The data was fitted with an exponential for the *in silico* (black dashed lines) and experimental results (red dashed lines). Color bar indicates diameter of apical and basal dendrites at different distances from soma. C. Predicted dendritic to somatic attenuation ratios for *in silico* EPSP amplitudes measured at different locations on apical (green dots) and basal dendrites (blue dots). Color code is similar to B. D. Somatic validation. For each injected somatic stimulus (the left most column, black) an exemplar voltage trace is plotted from the recorded data (middle column, green) and from the e-model’s response (the right most column, blue). E. Feature scores calculated based on the model responses and experimental recordings in D.

Next, to test whether the e-model could reproduce a wide range of somatic responses we used somatic validations. We aimed to compare the experimental data to the responses of the optimized model for stimuli that were not used in the course of optimizations. We considered three protocols (Ramp, sAHP and IDHyperpol, see Methods) (Fig.4D, first column). From the corresponding single-cell recordings (Fig.4D, second column) and model responses (Fig.4D, third column) we extracted a set of e-features and calculated their scores (Fig.4E). Most of the e-features of the e-model are less than five standard deviations away from the experimental e-features, except the ‘spike count’ and ‘time to first spike’ for the Ramp protocol (8 standard deviation instead).

To assess whether parameters of the L5PC e-model were well constrained by the e-features used in optimization we analyzed the parameter sensitivity of the e-model (Fig.5A). For this, we varied the value of one parameter at a time while keeping the rest of the parameters fixed. The values of the parameters were varied by 1%, 50% and 90%. Then, we computed the slope of the change in score value for each parameter (Fig.9). If the slope was approximately zero it meant that changes in the parameter value did not affect the e-feature in question. We demonstrate the results of this analysis for three protocols (bAP, Step_150% and IV_-100%). Our results show that the passive parameters (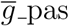, e_pas, 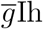), axonal sodium and somatic sodium demonstrated a slope larger than one for all e-features in all the protocols considered (Fig.5A). All axonal parameters, except transient potassium, somatic sodium and 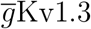 also show a slope larger than one for the e-features reflecting firing properties of the cell, such as interspike intervals and firing frequency (Fig.5A). The level of calcium at the apical point was affected by manipulation of maximum conductances of apical sodium, low threshold calcium, and calcium dynamics, while less by high threshold calcium channels (slope = 0.16). All e-features show low sensitivity to the following parameters: somatic and axonal 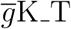 (maximum slopes are: 0.2 for the apical calcium amplitude; 0.16 for the time to last spike feature), somatic 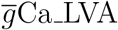 (slope: 0.14 for last AP amplitude) and apical 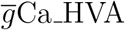 (slope 0.16 for feature: amplitude of calcium in apical dendrite) Fig.5A. These results indicate that the model is sensitive to changes in the majority of the parameters (26 out of 31).

**Figure 5:**
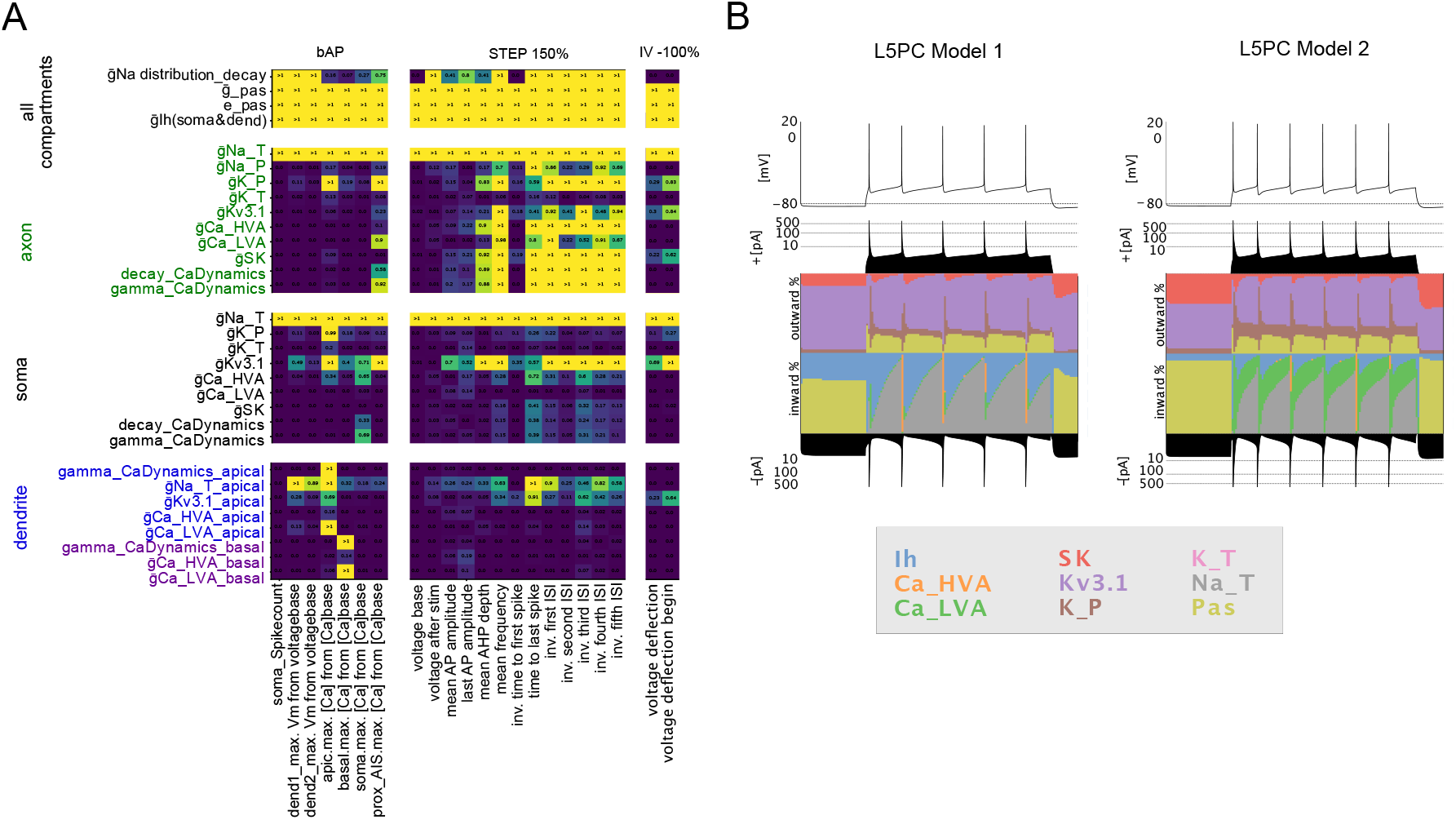
Sensitivity analysis and degeneracy. A. Analysis of the sensitivity of e-features to changes in the parameter values. The matrix represents slopes of e-features values. The sensitivity is presented for e-features extracted based on three protocols (bAP, Step 150% and IV —100%). Colors reflect the value of the slope, with slopes greater than one represented in yellow. B. Currentscape plots of two L5PC e-models, with two different sets of maximal intrinsic conductances. Top. Model responses to the same current stimuli (Step 150%). Bottom. Black-filled plots represent total positive (top) and negative (bottom) currents in the cell during the stimuli. The middle panel represents the contribution of ionic inward and outward currents during the stimuli, each color curve reveals the contribution of one particular ionic current as the percentage of the total current during the simulation.

Finally, we tested whether our modeling approach can reproduce the phenomenon of degeneracy of the ionic currents, that was previously reported to take place in electrical neuron models [Jain and Narayanan, 2020, Marder and Taylor, 2011, Rathour and Narayanan, 2019, Migliore et al., 2018, Drion et al., 2015, Goaillard and Marder, 2021]. For this purpose, we optimized another L5PC e-model, using different seeds for initialization of the e-model parameters. To illustrate the difference between the e-models we created currentscape plots [Alonso and Marder, 2019] of the somatic currents in response to the same percentage of injected current (depolarizing step of 150 % rheobase) (Fig.5B). We observed that the second e-model has a more prominent contribution of the Ca_LVA and less contribution of sodium current during firing than the first e-model. The difference between the e-models was also present in the contribution of potassium currents (K_P, Kv1.3, SK) and Ih. However, both e-models did not have any contribution of transient potassium (K_T) to the resting or firing states. This might indicate that K_T is not well constrained by the features used for optimization, independent of the optimization seed.

### 2.5 Generalization of electrical models

Each of the aforementioned optimized e-models was built for a particular morphology, which we call the exemplar morphology. When these e-models are meant to be used in a large network model, the question of the generalization of an e-model to other morphologies of the same or of different morphological type arises. Given the computational cost of optimizing an e-model and the possibly large number of reconstructed morphologies in the network, we chose to take the following approach. We assigned to each morphology the e-models that match it best, based on the most common e-types for a certain m-type.

An e-model for a morphology is accepted according to the rule in Equation 2, following the approach of Markram et al. [2015]. For this purpose we used a total of 1,015 reconstructed and manually corrected morphologies of SSCx young rat cortex for the generalization routine (see Methods). The population of morphologies was artificially increased to 141, 733 by a cloning procedure on the reconstructions [Reimann et al., 2022]. Given their m-type, morphologies were then attributed to one of the possible morpho-electro combinations [Markram et al., 2015, 2004] (see Methods), yielding 366,926 models, of which 233,941 passed generalization.

For the L5PC e-model we ran a generalization routine and separated morphologies on those that passed and failed this procedure (Fig.6A). We compared the distributions of e-feature scores between population of morphologies that failed and passed generalization (Fig.6B). We show that the e-features responsible for rejection of the morphologies are mostly related to firing properties and after-hyperpolarization depth.

**Figure 6:**
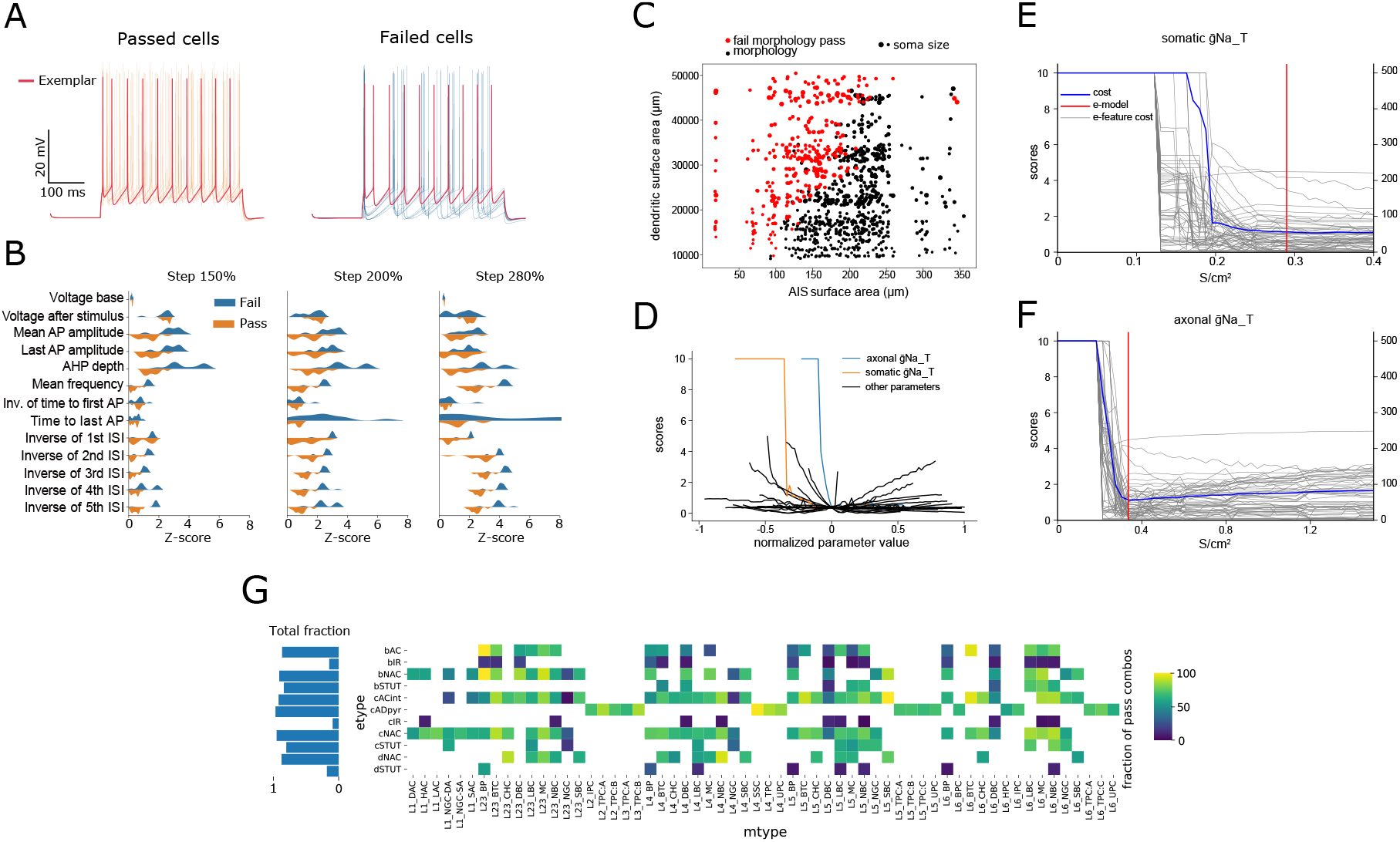
Generalization of electrical models. A. Examples of the traces for L5PC me-combination, in response to the depolarizing step of 150%. Traces of me-combinations that pass (left, orange and red traces) and (right, blue and red traces) the generalization procedure. Red traces corresponds to the optimized canonical e-model. B. Example of e-feature scores for me-combos that pass (orange) and fail (blue) generalization. E-features were extracted for three depolarization protocols (Steps 150%, 200%, 300%). C. Example of the morphological properties of L5PC me-combinations that passed and failed generalization. Plot illustrates relation between the total surface area of the AIS and (axon up to 40 *μm*) and the total surface area of the proximal dendritic compartments (up to 500 *μm* in path length for 1000 randomly sampled L5PC morphologies. Red dots represent failed morphologies, black dots for passed morphologies. Size of dots represents size of the soma. D. Parameter sensitivity analysis for the firing frequency e-feature in response to the depolarizing stimuli (150%). Dependency of e-feature score is plotted versus normalized value of the parameters. The blue line represents axonal sodium, orange for somatic sodium, black lines for all other parameters. Maximum score is clipped at 10. E-F. Sensitivity analysis with the exemplar morphology (similar to Fig.5) of the sodium conductance parameter in the axon (F) and soma (E) on the cost (red) and all features (gray). Red represents the value of these parameters for this cADpyr e-model (0.33 for axon and 0.29 for soma). G. For each combination of m-type and e-type, we display the fraction of accepted cells, with total fraction for each e-type on the left.

To gain a deeper insight into why some morphologies were not accepted for the L5PC emodel, we had a closer look at two morphological features (Fig.6C), the surface area of the axon initial segment (AIS) – taken here as the first 40 *μm* of axon – versus the proximal dendritic surface area (up to 500 *μm* in path length) for a random sample of 1000 morphologies. We observe a near linear correlation between these two quantities for a morphology to be valid under this e-model. These results are in line with previous works such as Hay et al. [2013] and Rall [1959], which have shown that a consistent *ρ* factor, defined as the ratio of input resistances between the AIS and somatodendritic compartments, is important to ensure generalizability of electrical models. Interestingly we noticed that soma surface area does not correlate with the pass/fail of the cell (Fig.6C).

Further we confirm that the sodium channels (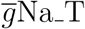 somatic and axonal) are indeed the most important for firing e-features such as spiking frequency (Fig.6D). These two sodium channels are able to bring the cell to a regime where threshold current is too high with respect to experimental recordings, thus no protocols can even be computed, resulting in maximum scores. For these two conductance parameters we show in panels (Fig.6E) (for soma) and (Fig.6F) (for axon) the sensitivity to all other e-features as well as the total cost (used for optimization) as a function of the non-normalized conductance values. We observe that the optimal solution is near the lowest possible 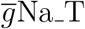 for the exemplar cell to fire, thus any small deviation of the ratio between the size of axon and the rest of the morphology may bring a cell into a non-firing regime. This transition point corresponds to the diagonal separation between black and red dots in panel (Fig.6C), as already pointed out in Hay et al. [2013]. On the contrary, the 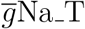 in the soma in Fig.6E is further away from the non-firing regime, thus the soma size is less important for a cell to be generalized on other morphologies. The optimized value of the axonal 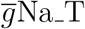 is close to the minimal value for the exemplar cell to be valid. Thus if a morphology with the same dendritic structure has an AIS with a smaller surface area, the cell will not pass our selection criteria.

With this morphological and parameter sensitivity analysis, we were able to determine that the low value of axonal 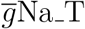 was the main cause of failed morphologies, suggesting that a slight manual adjustment of this parameter may help to improve the generalizability of this e-model.

Overall, our pipeline produces generalizable electrical models with a high acceptance rate for most e-types, except for three irregular electrical types: bIR, cIR and dSTUT (Fig.6G). For the dSTUT e-type only few me-models showed a delay in the first spikes (not shown), preventing most to pass. For cIR and bIR e-types, the spiking frequencies and interspikes interval are the most frequent e-features to fail, in addition to the burst number for bIR (meaning many me-models did not burst as expected) and the amplitudes of action potentials for cIR (not shown).

Finally, as a verification that the e-models obtained in this work are more generalizable than in a previous study (Markram et al. [2015]), we compare in Sect.5.1 the results of generalizability between both versions, and show a clear improvement using the present e-models.

## 3 Discussion

In this work we present an automatic workflow for single-cell model creation. We demonstrate the application of this workflow by producing 40 e-models for 11 e-types of the somatosensory cortex of a juvenile rat. These models reproduce neuronal responses from patch-clamp recordings observed in the corresponding cells. As an example of a single cell study, we assessed the L5PC e-model robustness, analyzed its parameters and tested the morphological generalization. We show that the optimized L5PC e-model can be successfully generalized to a wide range of L5PC morphologies and that the majority of the e-models parameters are constrained by the optimization cost function used in this study. Moreover, we show that L5PC models can reliably reproduce dendritic signal attenuation. These canonical e-models can be used to study signal propagation in a single cell or in networks. All the steps of the workflow are available to the reader as openly accessible Python notebooks (4.5).

The approach of building a canonical e-model is based on averaging neuronal responses across different cells that belong to the same e-type. Although we observe certain variability in the extracted e-features for each e-type (Fig.2), the canonical e-models will produce only a single e-feature value. This implies that variability present in the data is dismissed. This is not necessarily prohibitive in case one studies and analyzes average neuronal behavior, but can be problematic in tasks where variability is implied, such as circuit simulations. One direct way to overcome this limitation is to build e-models for every single neuron in the network. However, this would require a large amount of experimental data and be computationally expensive. It would also be possible to generate several e-models based on the same input, by varying the random generator seed, such that the resulting e-models have different optimized sets of parameters and therefore could represent variability present in the data. This would allow for the creation of several canonical e-models, each from the same population, yet allowing for the variability of neuronal responses. Alternatively, in this study we introduce variability among the me-models by generalizing the e-models over a large set of morphologies.

The set of e-models produced in this work is based on somatic patch clamp recordings and can be enriched with more details in future versions. For example, to allow for a study of neuronal synaptic and dendritic integration, the current models of L5PC would benefit from constraining the e-model by using dendritic calcium imaging data. Also several currents and channels could be included in the models, such as specific small and big conductance dendritic channels [Bock et al., 2019, Benhassine and Berger, 2005], or a nonlinear A-type K+ current distribution along the dendrites [Korngreen and Sakmann, 2000, Schaefer et al., 2007]. Moreover, by including axonal reconstructions and fitting corresponding axonal conductances, we could better understand their role in spike propagation and cell signaling.

In the present case of the SSCx interneuron optimizations we were able to successfully reproduce somatic responses of a wide range of e-types. For the irregular and stuttering interneurons, this was only possible by including stochastic potassium channels in the models. Interneuron e-models could be further refined through somatic validation analysis by quantifying the performance of optimized e-models on a battery of recordings that were not used in optimizations (via stimuli such as e.g. a ramp, a sinusoid, etc.). Finally, it should be mentioned that another approach to produce even more detailed e-models would be to incorporate transcriptomics data [Gouwens et al. [2018], Tasic et al. [2018], Nandi et al. [2020]], which would guide the composition of ionic currents present in the model. However, it is not clear yet how much the gene level expression is an indicator for the protein densities on the cellular membrane.

Yet another aspect of our approach to building a canonical e-model was that the optimization was performed on single morphology, while variability of morphologies is present within each m-type [Gouwens et al. [2019], Chen et al. [2009], Vrieler et al. [2019]]. We explored how well a single canonical e-model can be generalized on various morphologies of compatible m-types.

According to our results, while other interneurons had high total generalizability, irregularly firing and delayed stuttering interneurons showed poor generalizability. This could indicate that irregularly firing models are more sensitive than, for example, continuously firing e-types, to the morphological properties of the cells. We also observed that pyramidal cells show high generalizability. When we analyzed the morphological properties that allowed L5PC e-models to perform acceptably on a large set of morphologies, we noticed that the size of the AIS and the dendritic area play a considerable role in the e-model score. Most of the morphologies with a small AIS and a large dendritic area failed when applied to the canonical e-model, which is consistent with previously reported results [Hay et al., 2013]. To further understand and possibly improve the generalizability of canonical e-models one would need to study the link between morphological properties and electrical features of the e-models in more depth.

The parameters of the optimized e-model can be analyzed to assess the its biophysical properties. For example, with sensitivity analysis we showed how performance of the models is affected by the perturbations of the parameters [Jezzini et al., 2004, Tennøe et al., 2018]. In the case of the L5PC e-model, most of the parameters were constrained by the chosen evaluation function, while changes in some parameters such as axonal and somatic persistent potassium current maximal conductances had no effect on the e-features. This type of analysis may guide choices for e-features and protocols to consider in the score function, such that all parameters are sufficiently constrained. For a more in depth analysis, a closer look at the channel kinetics and its interplay with the rest of the currents in the cell may provide us more information about emodel sensitivity. Another aspect of the parameter space analysis is to look at the degeneracy of parameters [Marder and Taylor, 2011, Goaillard and Marder, 2021, Jain and Narayanan, 2020], meaning that we can produce several parameter sets which would result in similar performances of the e-model. We used currentscapes plots to visualize the current dynamics of two L5PC models with similar performance (reflected in the values of the evaluation function). We saw that ionic contributions to the overall current was different between the models. However, both models have a similar parameter sensitivity, with for example no contribution of persistent potassium current. Further, it could be interesting to investigate to what extent the definition of the score function can affect the presence of degeneracy in the system and to what extent it is an implicit property of the system.

## 4 Methods

### 4.1 Data

#### 4.1.1 Electrophysiological recordings

Electrophysiological recordings were obtained from P14-16 rat somatosensory cortex using whole-cell patch clamp experiments. The recordings were performed as described in Markram et al. [2015]. Each cell was classified according to its firing type based on the Petilla convention [Ascoli et al., 2008, Markram et al., 2015]. Recordings were performed in pyramidal cells (7 L6PCs, 44 L5PCs, 3 L4PCs, 8 L23PCs) and interneurons (16 cAC, 22 bAC, 19 cNAC, 28 bNAC, 7 dNAC, 11 dSTUT, 8 bSTUT, 10 cSTUT, 14 bIR, 6 cIR). A number of stimuli were applied for each cell: IDrest (depolarizing steps, sampling frequency: 10 kHz, duration: 2s), IDthresh (depolarizing steps, sampling frequency: 10 kHz, duration: 2 s), APWaveform (depolarizing steps, sampling frequency: 50 kHz, duration: 50 ms), IV (sequence of current steps, from hyperpolarization to depolarization, sampling frequency: 10 kHz, duration: 3 s), SpikeRec (sequence of brief depolarizing pairs with increasing interval, sampling frequency: 50 kHz, duration: 1.5 s), Ramp (ramp current, sampling frequency: 10 kHz, duration: 2s), sAHP (small depolarization currents, sampling frequency: 10 kHZ, duration: 2.5 s), IDHyperpol (depolarizing square pulses preceded by hyperpolarizing step, sampling frequency: 10 kHZ, duration: 3 s). For each cell a holding current was applied in order to keep offset voltage at −70 mV (before liquid junction potential correction of 14 mV).

#### 4.1.2 Morphologies

The following m-types were considered in this study:

- Inhibitory m-types: DAC: Descending Axon Cell; HAC: Horizontal Axon Cell; LAC:Large Axon Cell; NGC-DA: Neurogliaform Cell with dense axon; NGC: Neurogliaform Cell; NGC-SA: Neurogliaform Cell with sparse axon; SAC: Small Axon Cell; BP: Bipolar Cell; BTC: Bitufted Cell; CHC: Chandelier Cell; DBC: Double Bouquet Cell; LBC: Large Basket Cell; MC: Martinotti Cell; NBC: Nest Basket Cell; SBC: Small Basket Cell;
- Excitatory cell m-types: IPC: Inverted PC; BPC: Bipolar PC; HPC: Horizontal PC; TPC:A: Tufted PC, late bifurcation; TPC:B: Tufted PC, early bifurcation; TPC:C: Tufted PC, small tuft; UPC: Untufted PC; SSC: Spiny Stellate Cell.

A total of 1, 015 reconstructed rat morphologies ([Reimann et al., 2022, Ramaswamy et al., 2015], expanded dataset from Markram et al. [2015]) have been curated and repaired for cut plane missing data, following the procedure outlined in Markram et al. [2015]. In that study a cloning strategy was applied by successive application of rescaling operations and jitter of section lengths and bifurcations angles. This procedure was designed so that the resulting morphologies would retain their morphological types, while providing more variability across morphologies. In addition, axons and dendrites were shuffled within their original m-types to create more clones. We applied this procedure on the extended dataset of reconstructions to obtain a total of 141, 733 different morphologies. The relative proportions of morphological types is dictated by the expected density of morphologies of each type in SSCx, see Table 6.

### 4.2 Single cell model

#### 4.2.1 Electrophysiological features extraction

The extraction of e-features from the voltage traces was performed using the BluePyEfe python package [Blue Brain Project, 2020a]. A set of e-features (as described in sec 2.3 and sec 2.4) was extracted from each patch clamp recording for both optimizations (list of protocols and e-features Table 1) and validations (Table 2). The descriptions of the extracted features are summarized in Table 8. The membrane potential used as a detection threshold for the onset of an action potential is −30 mV. In addition, action potentials are only taken into consideration if they happen during a stimulus. The e-features are averaged across cells of the same e-type. The main issue with this process is that recordings coming from different cells have different input resistances. To solve this issue we first define “targets” expressed as amplitudes relative to the rheobase of the cells. The rheobase of each cell is computed as the lowest stimulus amplitude inducing a spike in any of its ‘IDrest’ or ‘IDthresh’ recordings. The stimulus amplitudes of all recordings were normalized by the rheobase of their respective cells. Finally, e-feature vectors were averaged across cells at the target levels. For this operation, a tolerance (10 % in the present study) is used as a binning width around the targets. For example, for a tolerance of 10 % and a target at 150 % rheobase, e-features are averaged for stimuli with amplitudes ranging from 140 to 160 %. The standard deviations of the e-features at the targets are computed following the same protocol.

**Table 1:**
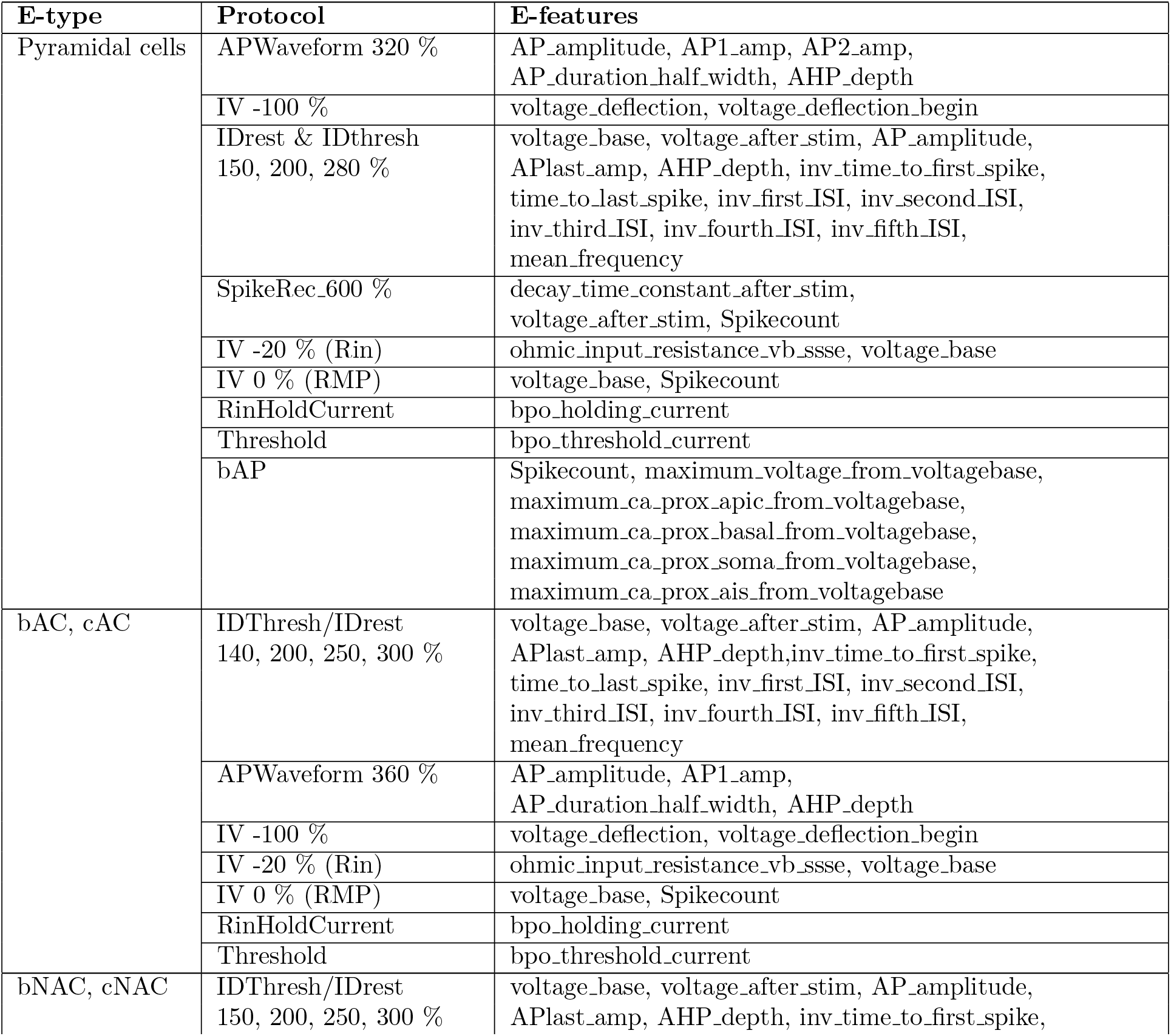
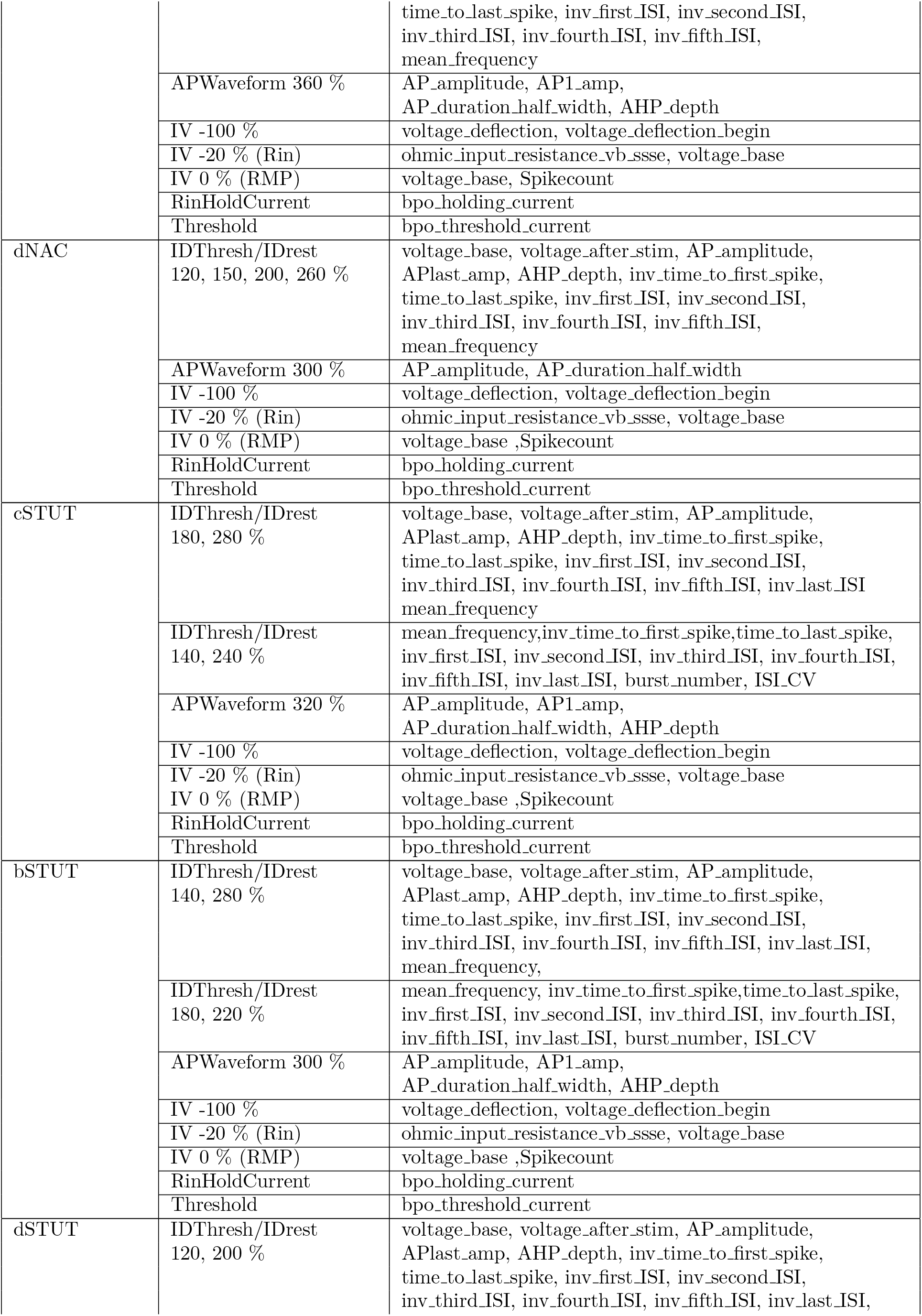
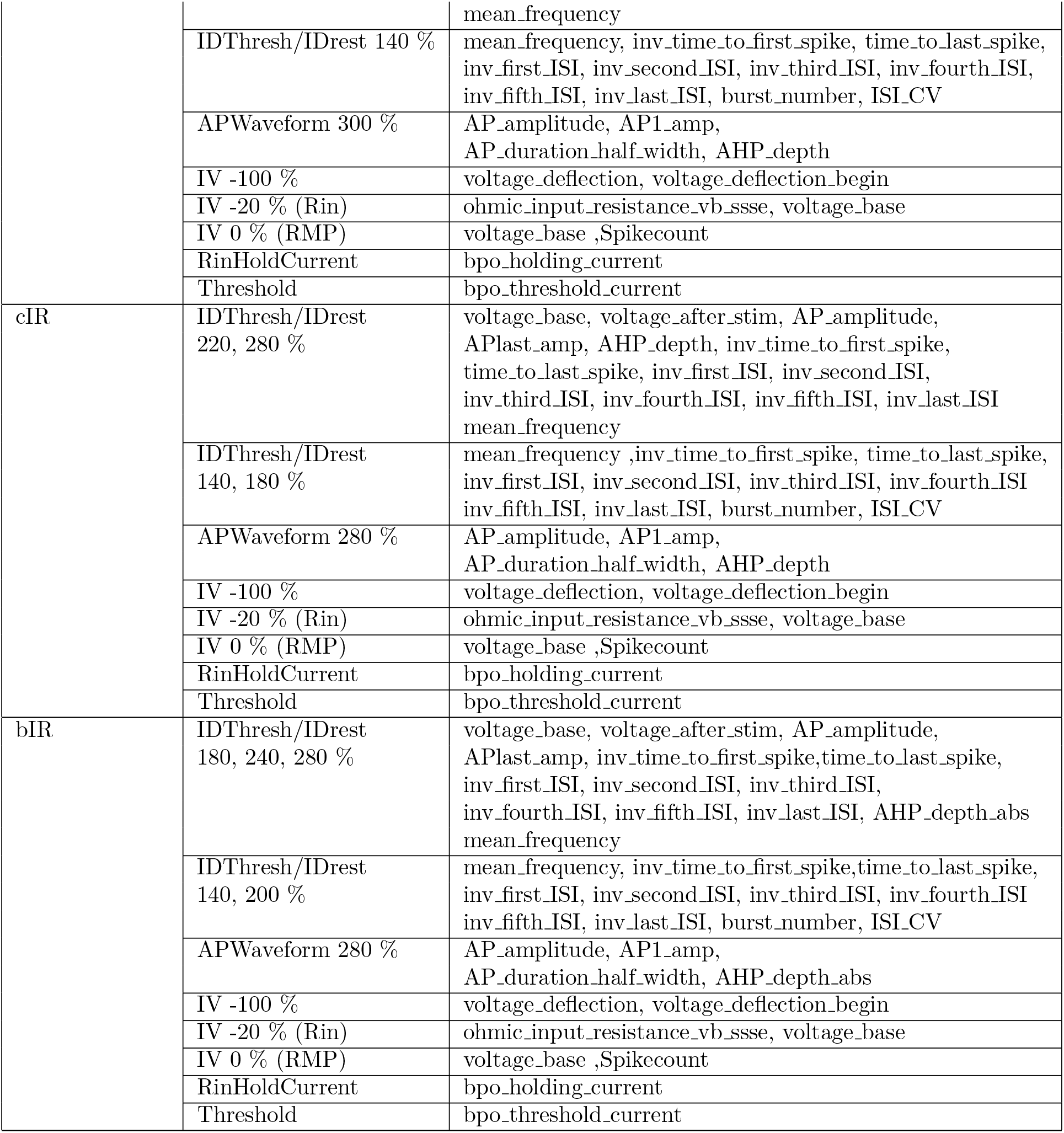
List of protocol and features used for optimizations for each e-type

#### 4.2.2 Single Neuron Models and Optimization

Morphologies were manually reconstructed. The axons and their branches were replaced by a synthetic axon section consisting of an AIS (60 *μ*m) followed by a myelinated axon segment of 1000 *μ*m.

The constant parameters used in the e-model are mentioned in Table 4. The mechanisms (ion channels and calcium dynamics) are listed in the Table 5. The various mechanisms added to neuron sections of the cADpyr e-type model is shown in Fig.3A and for other interneuron e-types listed in Table 7. The responses of the ion channels to step voltage clamp stimuli are plotted in Figure 7.

The optimization of the e-models was carried out using BluePyOpt Van Geit et al. [2016]. As evolutionary algorithm the IBEA [Zitzler and Künzli, 2004] method was chosen, with an offspring size of 256 individuals. E-features extracted from the experimental data as described above 4.2.1 were used as constraints. The protocols and features used for the optimizations are listed in Table 1.

We calculated the e-features scores using the formula:

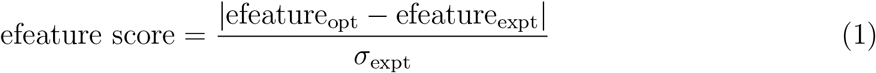

where e-feature_opt_ and e-feature_expt_ are the e-feature values from optimization and experiments, respectively, and σ_expt_ is the standard deviation of the experimental e-features.

During each evaluation of the model the following steps are followed:

1. Calculation of the resting membrane potential (RMP): For a set of e-model parameters obtained from the optimizer, we computed the soma RMP of the model when no stimulus is applied.
2. Calculation of the input resistance (Rin): We use a bisection search to find the holding current that brings the model to the same holding membrane potential as in the experimental Rin protocol.
3. Calculation of the threshold current: A bisection search is then also used to find the depolarizing threshold current for generating one spike in the model. If the scores for the Rin and RMP e-features are above given limits (3 standard deviation of respective mean e-feature value), the evaluation is stopped and no further protocols are applied.
4. Evaluation of the other protocols: If the model passes step 3, responses and e-features for other protocols such as IDRest, APWaveform and IV are evaluated.

The optimizations for the e-types cADpyr, bAC, cACint, bNAC, cNAC and dNAC were carried out in a single stage with the optimization algorithm runing for 100 generations (with an offspring size of 256). The models with irregular firing (bIR, cIR, bSTUT, cSTUT, and dSTUT) used a 2 stage optimization approach. These e-models include of stochastic potassium channel, StochKv3 to introduce the stochasticity in the firing patterns. The stochastic channel can work in two modes: deterministic and non-deterministic mode. In the first stage of the two staged optimization, all the variable e-model parameters including maximum conductance of stochastic channel (in deterministic mode) were optimized. In the second stage, all the optimized parameters from first stage are obtained and fixed except the stochastic channel conductances. The model is then optimized keeping non-deterministic mode on for this channel. The second stage of optimization was run for 50 generations with 256 offspring per generation.

#### 4.2.3 Model validation

Somatic and dendritic L5PC validations were performed using the BluePyEfe and BluePyOpt tools. The protocols and features used for somatic validations are listed in Table 2. In particular, several features were extracted in a specific manner:

- for the sAHP protocol: The “sag_ratio” and “sag_amplitude” were measured for the AHP that occurs after the short depolarization step.
- for the IDHyperpol protocol: The “sag_ratio” and “sag_amplitude” were measured during the hyperpolarizing step, while features that correspond to firing properties were measured during the depolarizing step.

**Table 2:** List of protocols and e-features used for validations of the L5PC model

| Protocol | E-features |
| --- | --- |
| IDHyperpol 250 % | Spikecount, AP_amplitude, ISI_values, ISI_log_slope, sag_amplitude, sag_ratio, minimum_voltage, steady_state_voltage_stimend |
| sAHP 150, 350 % | Spikecount, AP_amplitude, inv_time_to_first_spike, AHP_depth_abs, sag_ratio, decay_time_constant_after_stim, steady_state_voltage_stimend, sag_ratio, minimum_voltage, steady_state_voltage |
| APThresh 150 % | Spikecount, AP_amplitude, inv_first_ISI, AP1_amp, APlast_amp |

For the dendritic validations, *in silico bAP* recordings were performed with maximum distances from the soma of 900 μm for apical and 150 μm for basal dendrites. Diameters of apical dendrites were measured at the midpoint from soma to apical point, and for basal dendrites at the midpoint from soma to end point.

EPSP amplitude attenuation ratios were measured from the resting potential to the maximum of the EPSP during transient synaptic conductance change. These EPSPs were induced at each dendritic section measured at different locations on apical or basal dendrites. Exponential fits used the Levenberg-Marquardt algorithm [Levenberg, 1944].

### 4.3 Sensitivity analysis

The sensitivity analysis was performed for each parameter present in the e-model. Each parameter value was modified at a time, while the rest remained intact. At each step the e-feature values were computed. These values were compared with the control e-features (with all the parameter values remained intact). Each parameter was decreased by 10, 50 and 90 %. The final sensitivity value for the parameter was computed as a slope of the curve that represents the differences in e-feature between control and modified e-models:

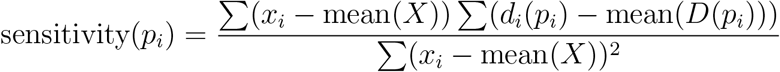

where *D*(*p_i_*) is e-features(control) – e-features(*p_i_*), *p_i_* the set of parameters where the ith parameter is modified, *X* corresponds to the percentage by which the ith parameter was modified.

### 4.4 Electrical model generalization

The procedure to assign e-models to each morphology to create me-models is as follows. For a given cloned morphology we selected a list of possible optimized e-models that match its me-type. We then evaluated the protocols used for the optimization and recorded the scores for each e-model.

We compared these scores with the scores computed on the exemplar morphology and accepted an me-model according to the following rule

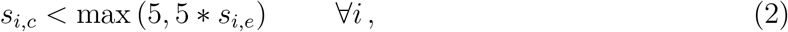

with *s_i,e_* the score of the exemplar morphology for a feature indexed by *i*, and *s_i,c_* the score of the me-model for the same feature. This procedure yielded 233’941 valid me-models which were then used to study the generalization of the e-models and to create a circuit model. The large number of evaluations was performed in parallel using the open source BluePyMM software [Blue Brain Project, 2020b].

### 4.5 Data and Code Availability

To illustrate the usage of our workflow we prepared a set of Python notebooks: https://github.com/BlueBrain/SSCxEModelExamples. They allow the reader to run each step of the pipeline: feature extraction, optimization, validation and generalization. The notebooks were developed for the L5PC example.

## 5 Supplementary Information

**Table 3:**
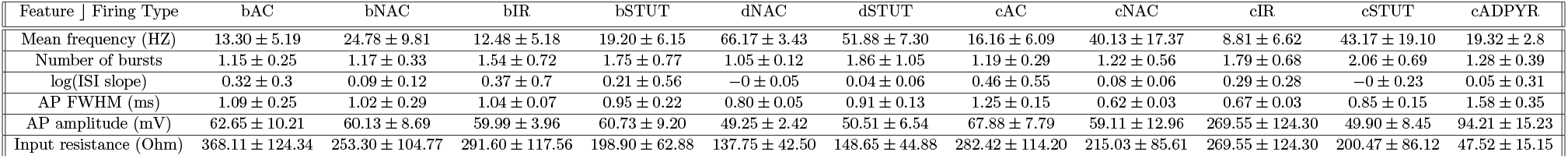
Values of the extracted e-features for 11 firing type as in Fig 2 B. The values are reported as mean ± SD

**Table 4:** Constant parameters of the electrical model

| Property | Value |
| --- | --- |
| Resting Membrane Potential (RMP) | -80 mV |
| Temperature | 34 °C |
| Specific Membrane Capacitance (cm), Soma | 1μF/cm<sup>2</sup> |
| Specific Membrane Capacitance (cm), Dendrites | 2μF/cm<sup>2</sup> |
| Specific Membrane Capacitance (cm), Myelinated Axon | 0.02μF/cm<sup>2</sup> |
| Axial Resistance | 100 Ωcm |
| Na<sup>+</sup> Nernst Potential | 50 mV |
| K<sup>+</sup> Nernst Potential | -90 mV |
| Initial Ca<sup>+</sup> Concentration | 6.5x10<sup>-5</sup> mM |

**Table 6:**
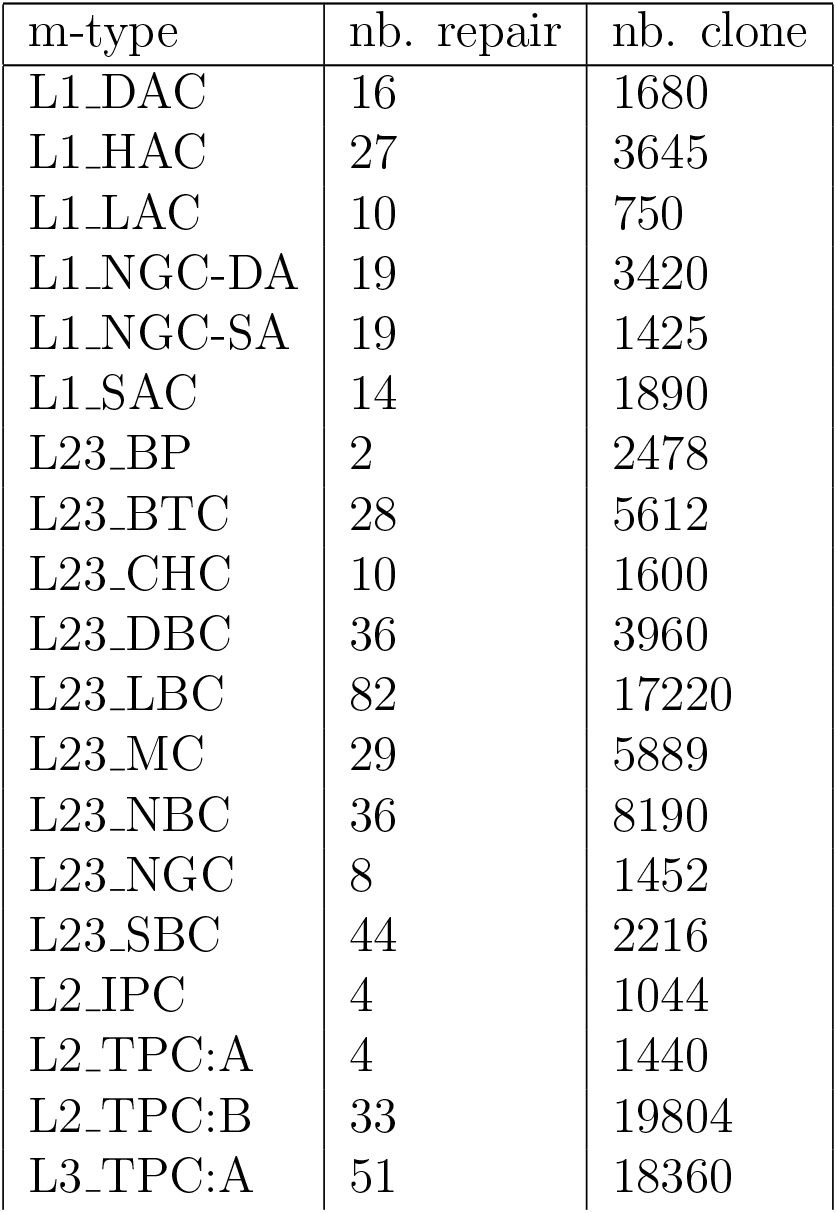
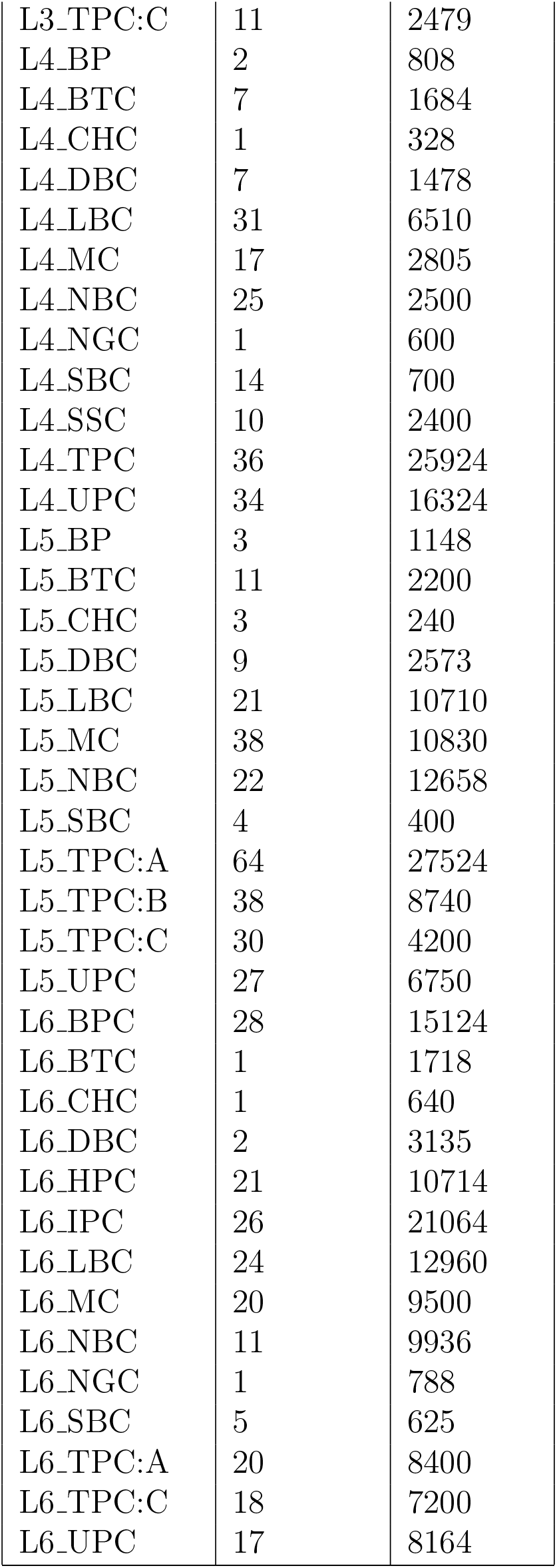
Number of repaired and cloned morphologies used for electrical model generalisation.

**Table 5:**
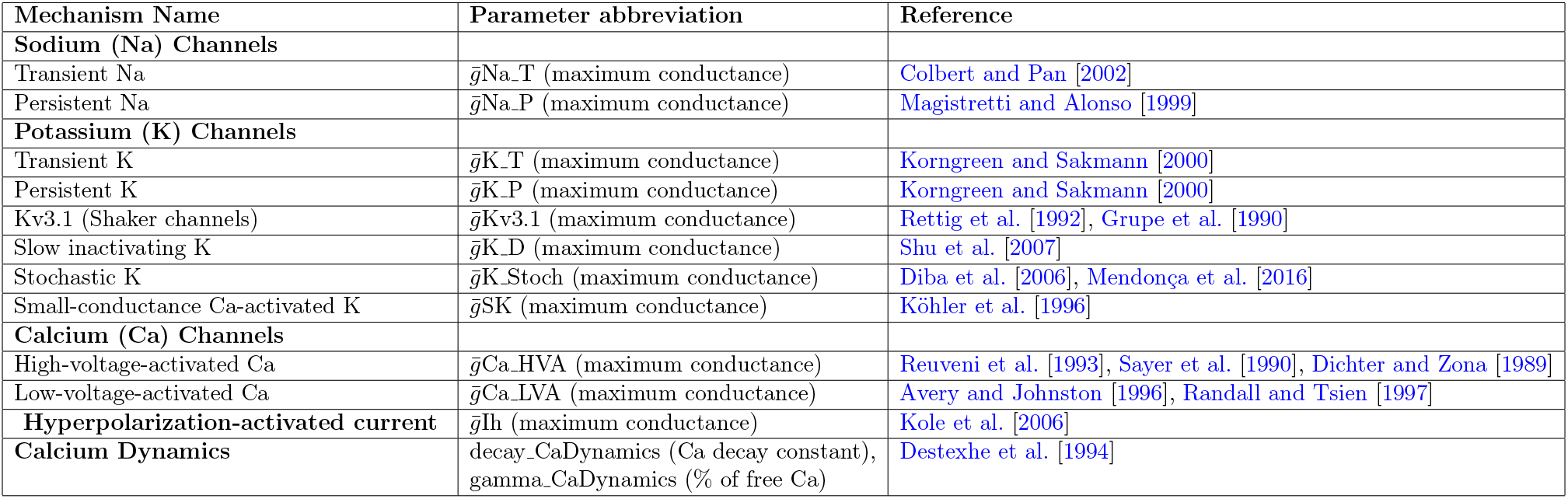
Active mechanisms and their respective parameters used for constructing e-models

**Figure 7:**
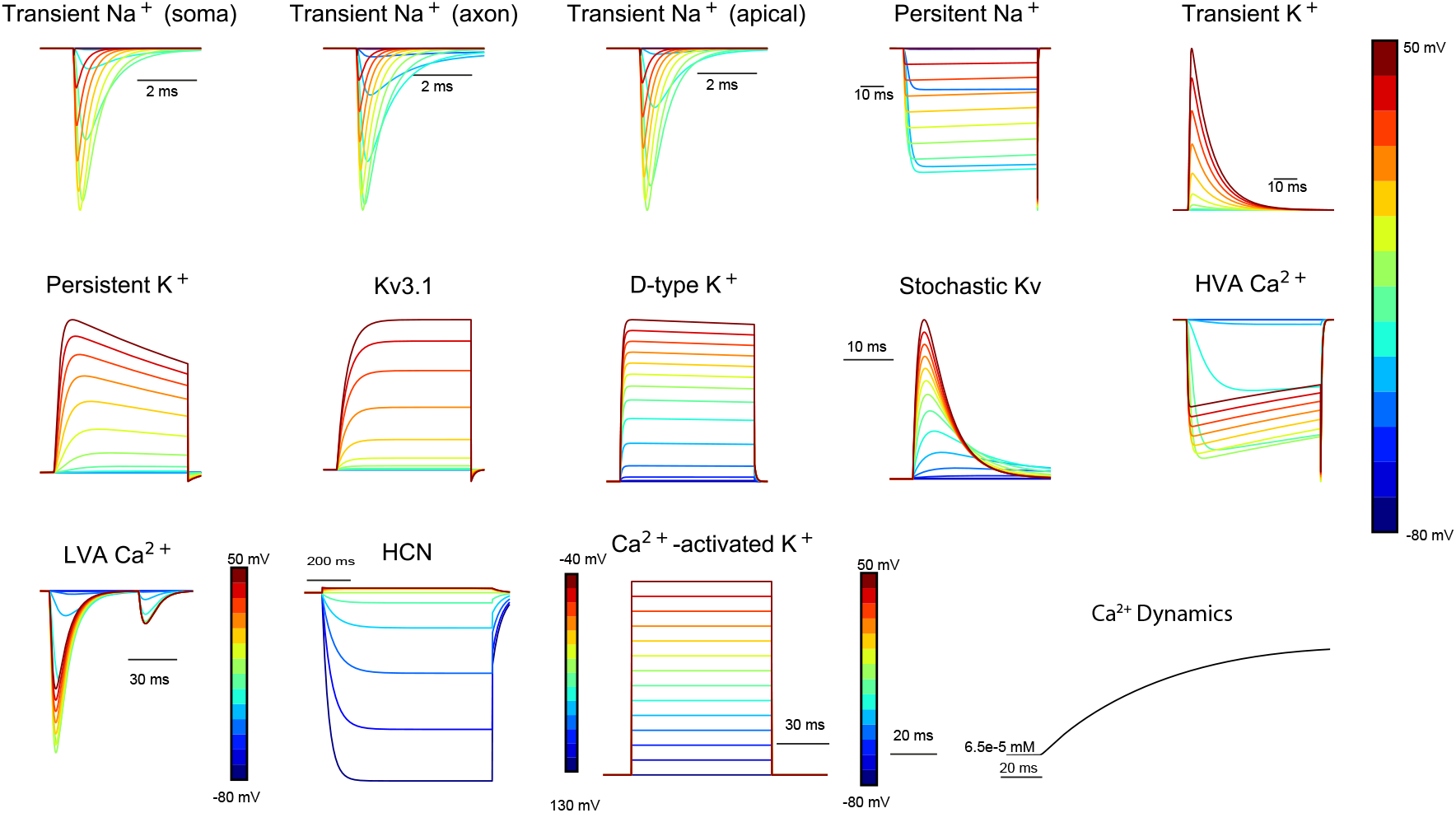
Ion channels currents used in cell models recorded for different voltage clamp injections. A calcium dynamics evolution with time is shown in black.

### 5.1 Comparison of generalization results between current and previous edition of the L5PC e-models

To perform generalization comparison we used the same set of e-features and same morphologies for two e-models: optimized in this work and previously reported L5PC e-model [Markram et al., 2015]. As a metric for this comparison we report the fraction of passed morphologies for each version of the e-models Fig.10.

The number of e-features computed in Markram et al. [2015] were smaller, without automatic holding and current threshold detection, and show a worse generalization ability on the morphologies used in this paper. Notice that these two e-models were trained on different exemplar morphologies, both present in the current set of morphologies.

**Table 7:** Recipe for each e-type: active parameters and their compartmental placement.

| E-type | Soma | AIS | Dendrites |
| --- | --- | --- | --- |
| cADpyr | CaDynamics<br>HVA Ca<br>LVA Ca<br>Kv3.1<br>Ca-activated K<br>Persistent K<br>Transient K<br>Transient Na<br>HCN | CaDynamics<br>HVA Ca<br>LVA Ca<br>Kv3.1<br>Ca-activated K<br>Persistent K<br>Transient K<br>Transient Na<br>Persistent Na | CaDynamics<br>HVA Ca<br>LVA Ca<br>Kv3.1 (Apical)<br>Transient Na (Apical; Decaying)<br>HCN (Apical and Basal;<br>Exponential increasing towards terminals) |
| bNAC, bAC, cNAC, cACint | CaDynamics<br>HVA Ca<br>LVA Ca<br>Ca-activated K<br>Transient Na<br>Kv3.1<br>Persistent K<br>Transient K<br>HCN | CaDynamics<br>HVA Ca<br>LVA Ca<br>Ca-activated K<br>Transient Na<br>Kv3.1<br>Persistent K<br>Transient K | CaDynamics<br>HVA Ca<br>LVA Ca<br>Ca-activated K<br>HCN (Exponential increasing towards terminals) |
| dNAC | CaDynamics<br>HVA Ca<br>LVA Ca<br>Ca-activated K<br>Transient Na<br>Kv3.1<br>Persistent K<br>Transient K<br>HCN<br>D-type K | CaDynamics<br>HVA Ca<br>LVA Ca<br>Ca-activated K<br>Transient Na<br>Kv3.1<br>Persistent K<br>Transient K<br>D-type K | CaDynamics<br>HVA Ca<br>LVA Ca<br>Ca-activated K<br>HCN (Exponential increasing towards terminals)<br>D-type K |
| bIR, cIR, bSTUT,cSTUT | Passive<br>CaDynamics<br>HVA Ca<br>LVA Ca<br>Ca-activated K<br>Transient Na<br>Kv3.1<br>Persistent K<br>Transient K<br>HCN<br>Stochastic Kv | Passive<br>CaDynamics<br>HVA Ca<br>LVA Ca<br>Ca-activated K<br>Transient Na<br>Kv3.1<br>Persistent K<br>Transient K<br>Stochastic Kv | Passive<br>CaDynamics<br>HVA Ca<br>LVA Ca<br>Ca-activated K<br>HCN (Exponential increasing towards terminals)<br>Stochastic Kv |
| dSTUT | CaDynamics<br>HVA Ca<br>LVA Ca<br>Ca-activated K<br>Transient Na<br>Kv3.1<br>Persistent K<br>Transient K<br>HCN<br>Stochastic Kv<br>D-type K | CaDynamics<br>HVA Ca<br>LVA Ca<br>Ca-activated K<br>Transient Na<br>Kv3.1<br>Persistent K<br>Transient K<br>Stochastic Kv<br>D-type K | CaDynamics<br>HVA Ca<br>LVA Ca<br>Ca-activated K<br>HCN (Exponential increasing towards terminals)<br>Stochastic Kv<br>D-type K |

**Table 8:** Descriptions of the eFEL feature used for optimizations and validations of emodels. More details for the features can be found at https://efel.readthedocs.io/en/latest/eFeatures.html.

| eFEL Feature | Description |
| --- | --- |
| Spikecount | Number of spikes in the trace, including outside of stimulus interval |
| maximum_voltage_from_voltagebase | Difference between maximum voltage during stimulus and voltage base |
| voltage_base | The average voltage during the last 10% of time before the stimulus. |
| voltage_after_stim | The mean voltage after the stimulus in (stim_end + 25%*end_period, stim_end + 75%*end_period) |
| AP_amplitude, APlast_amp, AP1_amp, AP2_amp | The relative height of the action potential from spike onset |
| AHP_depth | Relative voltage values at the first after-hyperpolarization |
| mean_frequency | The mean frequency of the firing rate |
| inv_time_to_first_spike | 1.0 over time to first spike; returns 0 when no spike |
| time_to_last_spike | time from stimulus start to last spike |
| inv_first_ISI, inv_second_ISI, inv_third_ISI, inv_fourth_ISI, inv_fifth_ISI, inv_last_ISI | 1.0 over first/second/third/fourth/fifth/last ISI; returns 0 when no ISI |
| AP_duration_half_width | Width of spike at half spike amplitude |
| ohmic_input_resistance_vb_ssse | The ratio between the voltage deflection (between voltage base and steady-state voltage at stimend) and stimulus current |
| voltage_deflection | The voltage deflection between voltage base and steady-state voltage at stimend |
| voltage_deflection_begin | The voltage deflection between voltage base and steady-state voltage soon after stimulation start. |
| decay_time_constant_after_stim | The decay time constant of the voltage right after the stimulus |
| holding_current | holding current injected to stabilize resting membrane potential |
| threshold_current | minimum current required to generate an AP found by bisection search |
| AHP_depth_abs | Absolute voltage values at the first after-hyperpolarization |
| burst_number | The number of bursts |
| ISI_CV | The coefficient of variation of the ISIs |
| time_to_first_spike | Time from the start of the stimulus to the maximum of the first peak |
| time_to_second_spike | Time from the start of the stimulus to the maximum of the second peak |

**Figure 8:**
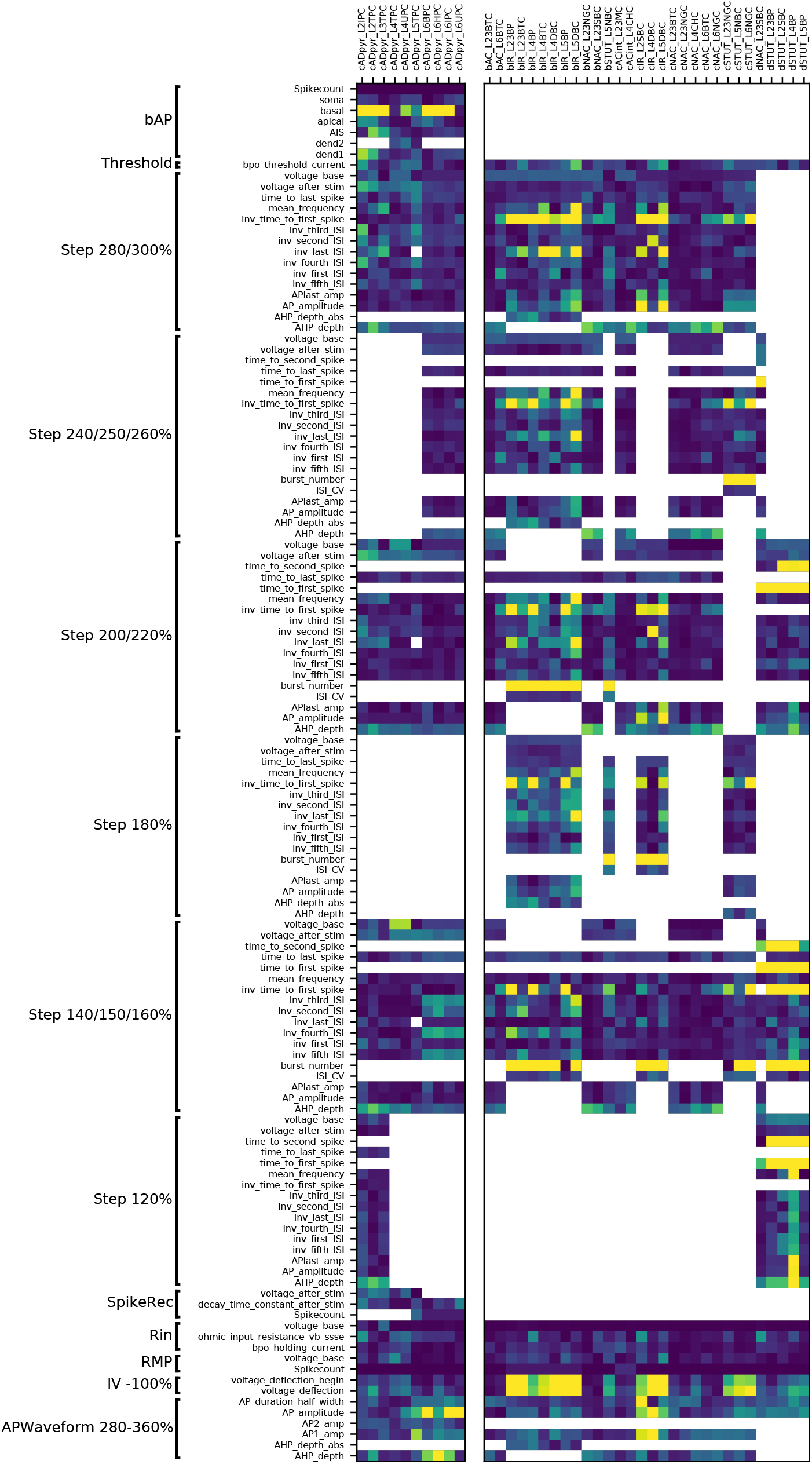
E-feature scores for all e-models optimized. E-feature descriptions can be found in Table 8. Blue represents a low and yellow a high score value, respectively.

**Figure 9:**
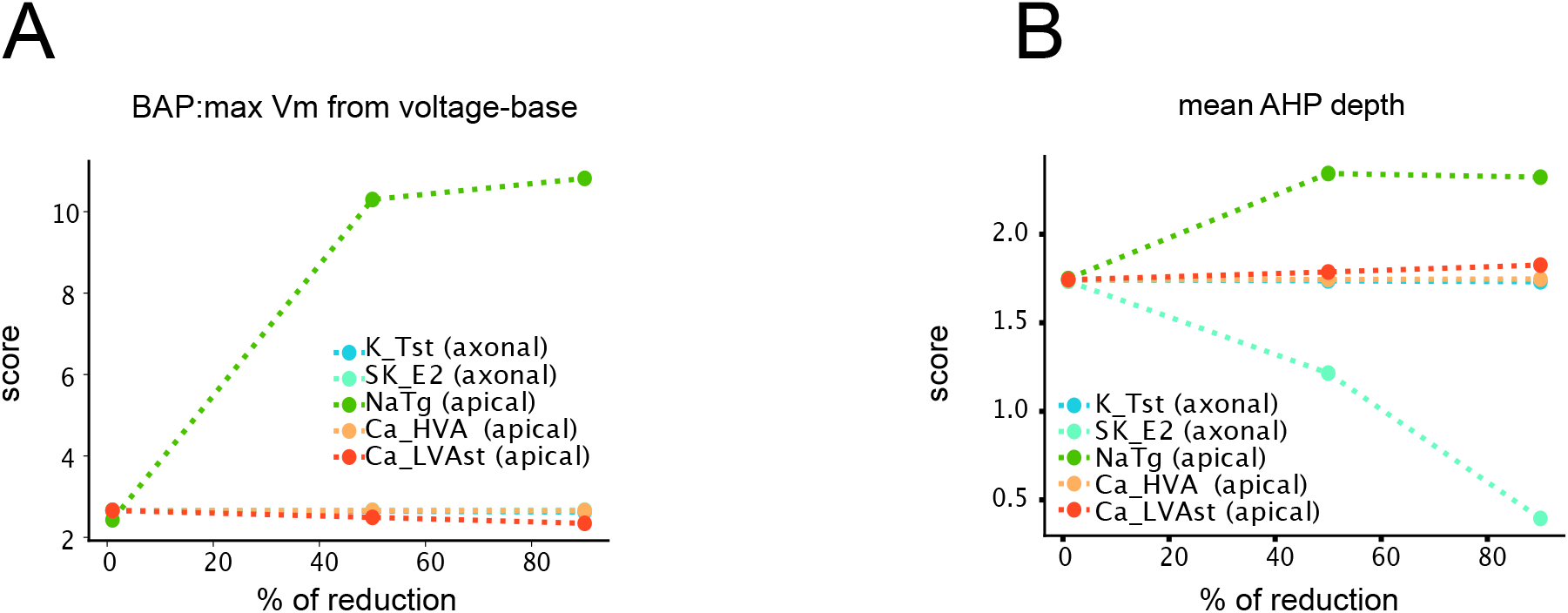
Examples of sensitivity analysis in L5PC models for the several parameters (K_Tst(axonal): blue, SK_E2(axonal): light blue, NaTg(apical): green, Ca_HVA(apical): orange, Ca_LVAst(apical): red), x-axis: percentage by which the parameter was reduced; y-axis: value of the e-feature score. A. Sensitivity analysis for the e-feature representing maximum voltage of the bAP. B. Sensitivity analysis for the mean AHP depth feature.

**Figure 10:**
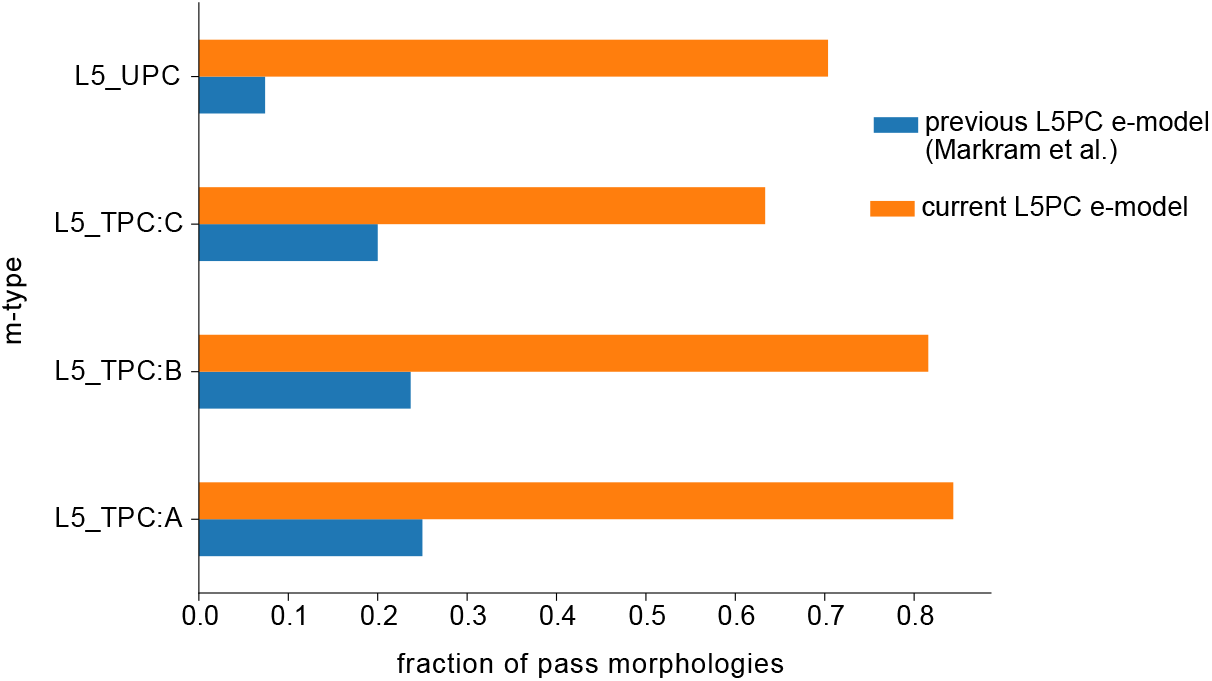
Comparison of generalization results between L5PC e-models from the current work (orange) and previously released e-model [Markram et al., 2015] (blue). Generalization was run for four morphological m-types of L5PC.

## References

L. M. Alonso and E. Marder. Visualization of currents in neural models with similar behavior and different conductance densities. Elife, 8:e42722, 2019.

G. A. Ascoli, L. Alonso-Nanclares, S. A. Anderson, G. Barrionuevo, R. Benavides-Piccione, A. Burkhalter, G. Buzsáki, B. Cauli, J. DeFelipe, A. Fairén, D. Feldmeyer, G. Fishell, Y. Fregnac, T. F. Freund, D. Gardner, E. P. Gardner, J. H. Goldberg, M. Helmstaedter, S. Hestrin, F. Karube, Z. F. Kisvárday, B. Lambolez, D. A. Lewis, O. Marin, H. Markram, A. Muñoz, A. Packer, C. C. H. Petersen, K. S. Rockland, J. Rossier, B. Rudy, P. Somogyi, J. F. Staiger, G. Tamas, A. M. Thomson, M. Toledo-Rodriguez, Y. Wang, D. C. West, R. Yuste, and T. P. I. N. G. (PING). Petilla terminology: nomenclature of features of gabaergic interneurons of the cerebral cortex. Nature Reviews Neuroscience, 9(7):557–568, 2008.

R. B. Avery and D. Johnston. Multiple channel types contribute to the low-voltage-activated calcium current in hippocampal ca3 pyramidal neurons. Journal of Neuroscience, 16(18): 5567–5582, 1996.

N. Benhassine and T. Berger. Homogeneous distribution of large-conductance calciumdependent potassium channels on soma and apical dendrite of rat neocortical layer 5 pyramidal neurons. European Journal of Neuroscience, 21(4):914–926, 2005.

T. Berger, M. E. Larkum, and H.-R. Lüscher. High i h channel density in the distal apical dendrite of layer v pyramidal cells increases bidirectional attenuation of epsps. Journal of neurophysiology, 85(2):855–868, 2001.

Y. N. Billeh, B. Cai, S. L. Gratiy, K. Dai, R. Iyer, N. W. Gouwens, R. Abbasi-Asl, X. Jia, J. H. Siegle, S. R. Olsen, et al. Systematic integration of structural and functional data into multi-scale models of mouse primary visual cortex. Neuron, 106(3):388–403, 2020.

Blue Brain Project. Blue brain python e-feature extraction library (bluepyefe). https://github.com/BlueBrain/BluePyEfe, 2020a.

Blue Brain Project. Blue brain python cell model management (bluepymm). https://github.com/BlueBrain/BluePyMM, 2020b.

T. Bock, S. Honnuraiah, and G. J. Stuart. Paradoxical excitatory impact of sk channels on dendritic excitability. Journal of Neuroscience, 39(40):7826–7839, 2019.

T. Brookings, M. L. Goeritz, and E. Marder. Automatic parameter estimation of multicompartmental neuron models via minimization of trace error with control adjustment. Journal of neurophysiology, 112(9):2332–2348, 2014.

N. T. Carnevale and M. L. Hines. The NEURON book. Cambridge University Press, 2006.

C.-C. Chen, S. Abrams, A. Pinhas, and J. C. Brumberg. Morphological heterogeneity of layer vi neurons in mouse barrel cortex. Journal of Comparative Neurology, 512(6):726–746, 2009.

C. M. Colbert and E. Pan. Ion channel properties underlying axonal action potential initiation in pyramidal neurons. Nature neuroscience, 5(6):533–538, 2002.

A. Destexhe, D. Contreras, T. J. Sejnowski, and M. Steriade. A model of spindle rhythmicity in the isolated thalamic reticular nucleus. Journal of neurophysiology, 72(2):803–818, 1994.

K. Diba, C. Koch, and I. Segev. Spike propagation in dendrites with stochastic ion channels. Journal of computational neuroscience, 20(1):77–84, 2006.

M. A. Dichter and C. Zona. Calcium currents in cultured rat cortical neurons. Brain research, 492(1-2):219–229, 1989.

G. Drion, T. O’Leary, and E. Marder. Ion channel degeneracy enables robust and tunable neuronal firing rates. Proceedings of the National Academy of Sciences, 112(38):E5361–E5370, 2015.

J.-M. Goaillard and E. Marder. Ion channel degeneracy, variability, and covariation in neuron and circuit resilience. Annual review of neuroscience, 44, 2021.

N. W. Gouwens, J. Berg, D. Feng, S. A. Sorensen, H. Zeng, M. J. Hawrylycz, C. Koch, and A. Arkhipov. Systematic generation of biophysically detailed models for diverse cortical neuron types. Nature communications, 9(1):1–13, 2018.

N. W. Gouwens, S. A. Sorensen, J. Berg, C. Lee, T. Jarsky, J. Ting, S. M. Sunkin, D. Feng, C. A. Anastassiou, E. Barkan, et al. Classification of electrophysiological and morphological neuron types in the mouse visual cortex. Nature neuroscience, 22(7):1182–1195, 2019.

A. Grupe, K. Schröter, J. Ruppersberg, M. Stocker, T. Drewes, S. Beckh, and O. Pongs. Cloning and expression of a human voltage-gated potassium channel. a novel member of the rck potassium channel family. The EMBO journal, 9(6):1749–1756, 1990.

C. Günay, J. R. Edgerton, and D. Jaeger. Channel density distributions explain spiking variability in the globus pallidus: a combined physiology and computer simulation database approach. Journal of Neuroscience, 28(30):7476–7491, 2008.

E. Hay, S. Hill, F. Schürmann, H. Markram, and I. Segev. Models of neocortical layer 5b pyramidal cells capturing a wide range of dendritic and perisomatic active properties. PLoS computational biology, 7(7):e1002107, 2011.

E. Hay, F. Schürmann, H. Markram, and I. Segev. Preserving axosomatic spiking features despite diverse dendritic morphology. Journal of neurophysiology, 109(12):2972–2981, 2013.

A. Jain and R. Narayanan. Degeneracy in the emergence of spike-triggered average of hippocampal pyramidal neurons. Scientific reports, 10(1):1–14, 2020.

S. H. Jezzini, A. A. Hill, P. Kuzyk, and R. L. Calabrese. Detailed model of intersegmental coordination in the timing network of the leech heartbeat central pattern generator. Journal of neurophysiology, 91(2):958–977, 2004.

M. Köhler, B. Hirschberg, C. Bond, J. M. Kinzie, N. Marrion, J. Maylie, and J. Adelman. Small-conductance, calcium-activated potassium channels from mammalian brain. Science, 273(5282):1709–1714, 1996.

M. H. Kole, S. Hallermann, and G. J. Stuart. Single ih channels in pyramidal neuron dendrites: properties, distribution, and impact on action potential output. Journal of Neuroscience, 26 (6):1677–1687, 2006.

A. Korngreen and B. Sakmann. Voltage-gated k+ channels in layer 5 neocortical pyramidal neurones from young rats: subtypes and gradients. The Journal of physiology, 525(3):621–639, 2000.

K. Levenberg. A method for the solution of certain non-linear problems in least squares. Quarterly of applied mathematics, 2(2):164–168, 1944.

J. Magistretti and A. Alonso. Biophysical properties and slow voltage-dependent inactivation of a sustained sodium current in entorhinal cortex layer-ii principal neurons: a whole-cell and single-channel study. The Journal of general physiology, 114(4):491–509, 1999.

Z. F. Mainen and T. J. Sejnowski. Influence of dendritic structure on firing pattern in model neocortical neurons. Nature, 382(6589):363–366, 1996.

E. Marder and A. L. Taylor. Multiple models to capture the variability in biological neurons and networks. Nature neuroscience, 14(2):133–138, 2011.

H. Markram, M. Toledo-Rodriguez, Y. Wang, A. Gupta, G. Silberberg, and C. Wu. Interneurons of the neocortical inhibitory system. Nature reviews neuroscience, 5(10):793–807, 2004.

H. Markram, E. Muller, S. Ramaswamy, M. W. Reimann, M. Abdellah, C. A. Sanchez, A. Ailamaki, L. Alonso-Nanclares, N. Antille, S. Arsever, et al. Reconstruction and simulation of neocortical microcircuitry. Cell, 163(2):456–492, 2015.

P. R. Mendonça, M. Vargas-Caballero, F. Erdélyi, G. Szabó, O. Paulsen, and H. P. Robinson. Stochastic and deterministic dynamics of intrinsically irregular firing in cortical inhibitory interneurons. Elife, 5:e16475, 2016.

R. Migliore, C. A. Lupascu, L. L. Bologna, A. Romani, J.-D. Courcol, S. Antonel, W. A. Van Geit, A. M. Thomson, A. Mercer, S. Lange, et al. The physiological variability of channel density in hippocampal ca1 pyramidal cells and interneurons explored using a unified data-driven modeling workflow. PLoS computational biology, 14(9):e1006423, 2018.

M. Mohácsi, M. P. Török, S. Sáray, and S. Káli. A unified framework for the application and evaluation of different methods for neural parameter optimization. In 2020 International Joint Conference on Neural Networks (IJCNN), pages 1–7. IEEE, 2020.

A. Nandi, T. Chartrand, W. V. Geit, A. Buchin, Z. Yao, S. Y. Lee, Y. Wei, B. Kalmbach, B. Lee, E. Lein, J. Berg, U. Sümbül, C. Koch, B. Tasic, and C. A. Anastassiou. Single-neuron models linking electrophysiology, morphology and transcriptomics across cortical cell types. bioRxiv, 2020. doi: 10.1101/2020.04.09.030239.

T. Nevian, M. E. Larkum, A. Polsky, and J. Schiller. Properties of basal dendrites of layer 5 pyramidal neurons: a direct patch-clamp recording study. Nature neuroscience, 10(2): 206–214, 2007.

P. Poirazi, T. Brannon, and B. W. Mel. Arithmetic of subthreshold synaptic summation in a model ca1 pyramidal cell. Neuron, 37(6):977–987, 2003.

A. A. Prinz, C. P. Billimoria, and E. Marder. Alternative to hand-tuning conductance-based models: construction and analysis of databases of model neurons. Journal of neurophysiology, 2003.

W. Rall. Branching dendritic trees and motoneuron membrane resistivity. Experimental neurology, 1(5):491–527, 1959.

S. Ramaswamy, J.-D. Courcol, M. Abdellah, S. R. Adaszewski, N. Antille, S. Arsever, G. Atenekeng, A. Bilgili, Y. Brukau, A. Chalimourda, et al. The neocortical microcircuit collaboration portal: a resource for rat somatosensory cortex. Frontiers in neural circuits, 9: 44, 2015.

A. Randall and R. Tsien. Contrasting biophysical and pharmacological properties of t-type and r-type calcium channels. Neuropharmacology, 36(7):879–893, 1997.

R. K. Rathour and R. Narayanan. Degeneracy in hippocampal physiology and plasticity. Hippocampus, 29(10):980–1022, 2019.

M. W. Reimann, S. B. Puchet, D. E. Santander, J.-D. Courcol, A. Arnaudon, B. Coste, T. Delemontex, A. Devresse, H. Dictus, A. Dietz, et al. Modeling and simulation of rat non-barrel somatosensory cortex. part i: Modeling anatomy. bioRxiv, 2022.

J. Rettig, F. Wunder, M. Stocker, R. Lichtinghagen, F. Mastiaux, S. Beckh, W. Kues, P. Pedarzani, K. H. Schröter, and J. P. Ruppersberg. Characterization of a shaw-related potassium channel family in rat brain. The EMBO Journal, 11(7):2473–2486, 1992.

I. Reuveni, A. Friedman, Y. Amitai, and M. J. Gutnick. Stepwise repolarization from ca2+ plateaus in neocortical pyramidal cells: evidence for nonhomogeneous distribution of hva ca2+ channels in dendrites. Journal of Neuroscience, 13(11):4609–4621, 1993.

R. Sayer, P. Schwindt, and W. Crill. High-and low-threshold calcium currents in neurons acutely isolated from rat sensorimotor cortex. Neuroscience letters, 120(2):175–178, 1990.

A. T. Schaefer, M. Helmstaedter, A. C. Schmitt, D. Bar-Yehuda, M. Almog, H. Ben-Porat, B. Sakmann, and A. Korngreen. Dendritic voltage-gated k+ conductance gradient in pyramidal neurones of neocortical layer 5b from rats. The Journal of physiology, 579(3):737–752, 2007.

I. Segev and M. London. Untangling dendrites with quantitative models. Science, 290(5492): 744–750, 2000.

V. Sekulić, J. J. Lawrence, and F. K. Skinner. Using multi-compartment ensemble modeling as an investigative tool of spatially distributed biophysical balances: application to hippocampal oriens-lacunosum/moleculare (o-lm) cells. PloS one, 9(10):e106567, 2014.

Y. Shu, Y. Yu, J. Yang, and D. A. McCormick. Selective control of cortical axonal spikes by a slowly inactivating k+ current. Proceedings of the National Academy of Sciences, 104(27): 11453–11458, 2007.

B. Tasic, Z. Yao, L. T. Graybuck, K. A. Smith, T. N. Nguyen, D. Bertagnolli, J. Goldy, E. Garren, M. N. Economo, S. Viswanathan, et al. Shared and distinct transcriptomic cell types across neocortical areas. Nature, 563(7729):72–78, 2018.

S. Tennøe, G. Halnes, and G. T. Einevoll. Uncertainpy: a python toolbox for uncertainty quantification and sensitivity analysis in computational neuroscience. Frontiers in neuroinformatics, 12:49, 2018.

R. D. Traub, E. H. Buhl, T. Gloveli, and M. A. Whittington. Fast rhythmic bursting can be induced in layer 2/3 cortical neurons by enhancing persistent na+ conductance or by blocking bk channels. Journal of neurophysiology, 89(2):909–921, 2003.

R. A. van Elburg and A. van Ooyen. Impact of dendritic size and dendritic topology on burst firing in pyramidal cells. PLoS computational biology, 6(5):e1000781, 2010.

W. Van Geit, E. De Schutter, and P. Achard. Automated neuron model optimization techniques: a review. Biological cybernetics, 99(4):241–251, 2008.

W. Van Geit, M. Gevaert, G. Chindemi, C. Rössert, J.-D. Courcol, E. B. Muller, F. Schürmann, I. Segev, and H. Markram. Bluepyopt: Leveraging open source software and cloud infrastructure to optimise model parameters in neuroscience. Frontiers in Neuroinformatics, 10 (17), 2016. ISSN 1662-5196. doi: 10.3389/fninf.2016.00017.

P. Vetter, A. Roth, and M. Häusser. Propagation of action potentials in dendrites depends on dendritic morphology. Journal of neurophysiology, 85(2):926–937, 2001.

N. Vrieler, S. Loyola, Y. Yarden-Rabinowitz, J. Hoogendorp, N. Medvedev, T. M. Hoogland, C. I. De Zeeuw, E. De Schutter, Y. Yarom, M. Negrello, et al. Variability and directionality of inferior olive neuron dendrites revealed by detailed 3d characterization of an extensive morphological library. Brain Structure and Function, 224(4):1677–1695, 2019.

J. J. Zhu. Maturation of layer 5 neocortical pyramidal neurons: amplifying salient layer 1 and layer 4 inputs by ca2+ action potentials in adult rat tuft dendrites. The Journal of physiology, 526(3):571–587, 2000.

E. Zitzler and S. Künzli. Indicator-based selection in multiobjective search. In International conference on parallel problem solving from nature, pages 832–842. Springer, 2004.

